# The genetic architecture of polygenic adaptation under a network-derived trait

**DOI:** 10.1101/2025.02.20.639381

**Authors:** Nicholas L. V. O’Brien, Barbara Holland, Jan Engelstädter, Daniel Ortiz-Barrientos

## Abstract

The genetic architecture of adaptation varies across species, populations, and traits. While existing models capture aspects like the number of loci, effect sizes, and allele frequencies, they often overlook the molecular processes underlying trait expression. We investigated how gene regulatory networks influence quantitative variation during adaptation by examining the negative autoregulation (NAR) motif in two configurations: K+, with four evolving network components, and K-, with two components. Using forward-time simulations, we tracked populations adapting to a shifted phenotypic optimum under varying genetic architectures. We found that K+ populations maintained rapid adaptation despite low recombination, preserving high genetic variance through positive epistasis and stronger linkage disequilibrium. Under low recombination, K+ populations reached the optimum through diverse molecular configurations, while responses were more uniform under high recombination. In contrast, K− and additive models showed impaired adaptation under low recombination. Our findings demonstrate that network structure fundamentally influences the distribution of variation within the molecular architecture of traits, with certain networks providing unexpected robustness against low recombination rates. This suggests that molecular complexity may confer evolutionary advantages in natural populations.

## Introduction

The ability of natural populations to adapt to changing environments is highly variable. Environmental changes bring about selective pressures, which might be insurmountable if the population lacks variation in traits under selection (Fisher 1930). Hence, adaptability is a function of *which* traits are under selection and how the genetic bases of those traits permit the production of variation that is a) congruous with the environment and b) reliably inherited from parent to offspring (i.e., “additive” in nature; Draghi and Whitlock 2012; Barghi *et al*. 2020). Because certain genetic architectures facilitate adaptation better than others, understanding the connection between genotype and phenotype is of great interest (Barghi *et al*. 2020).

The genetic architecture of a trait includes allelic effect sizes and frequencies, the number of causal loci, and recombination rates (Mackay 2001; Visscher and Goddard 2019). Quantitative traits often have genetic architectures comparable to Fisher’s infinitesimal model where many independent quantitative trait loci (QTLs) contribute small amounts of additive variation to the trait (Fisher 1918). The classic example is human height, where over 100,000 SNPs contribute to trait variation (Boyle *et al*. 2017), and approximately 80% of phenotypic variance is explained by additive genetic effects (Macgregor *et al*. 2006; Visscher *et al*. 2008). However, this additive framework may not fully capture the complexity of adaptive traits. Often, life history traits are subject to genetic interactions (Burch *et al*. 2024), suggesting that nonlinear genetic architectures might be necessary for driving variation in traits relevant to fitness.

Genetic interactions emerge from the molecular machinery underlying trait expression - specifically, the gene regulatory networks that orchestrate gene expression and trait development. Such networks create a hierarchical structure where genetic effects cascade through multiple regulatory layers (Claringbould *et al*. 2017; Fagny and Austerlitz 2021), transforming the simple additive model into a more complex system of interacting components (Nijhout and Paulsen 1997) dependent on their genetic background. For example, life history traits frequently display genetic interactions that cannot be explained by additive effects alone (Burch *et al*. 2024), suggesting that their variation is driven by the intricate relationships between regulatory elements within these networks. Furthermore, because the genetic background becomes an important component of an allele’s fitness, processes that control the co-inheritance of alleles, such as recombination, should be a key component of adaptation when epistasis is common (Otto 2009).

The evolution of recombination has long been associated with epistasis via its effects on linkage disequilibrium, LD (Otto 2009; Barton and Charlesworth 1998). LD measures the relative frequencies of genotypes, identifying whether certain combinations of alleles at different loci are inherited more or less often than expected from their frequencies in the population (Slatkin 2008). Epistasis affects LD depending on its sign: when epistasis is positive, combinations of mutations have greater fitness than expected from their individual effects. When epistasis is negative, combinations have lower fitness than expected. Positive epistasis tends to generate positive LD as selection maintains beneficial combinations, while negative epistasis leads to negative LD as selection favors breaking up disadvantageous combinations (Otto 2009; Barton 1995).

As LD accrues, changes in the genetic variance in fitness can manifest. Recombination reduces LD over time, which may increase or decrease variance in fitness, depending on the sign of LD. When LD is positive, genetic variance is maximized because extreme fitness genotypes are common in the population. The opposite is true when LD is negative (Eshel and Feldman 1970; Barton 1995). Given the ramifications for LD on fitness, understanding how a trait’s genetic architecture contributes to generating epistasis (and how recombination might interplay with said architectures) is a fundamental question in evolutionary genetics.

Gene regulatory networks are ideal for studying these relationships, as they naturally generate epistasis through their inherent complexity. Understanding how network architecture influences epistasis and, consequently, the maintenance of genetic variation is crucial for predicting adaptive trajectories in natural populations. Here, we investigate these dynamics using the negative autoregulation (NAR) motif, a ubiquitous subnetwork found in upwards of 40% of all *Escherichia coli* transcription networks (Alon 2007). The NAR network controls the expression of a gene *Z*, which serves as our quantitative trait. In our model (previously outlined in O’Brien *et al*. 2024), loci contribute to “molecular components” that modulate the shape of the *Z* expression curve, allowing genetic interactions to emerge naturally during evolution.

Using forward-time simulations, we examine how network structure influences adaptation under varying recombination rates and genetic architectures. By analyzing the distribution of additive variance across molecular components and the emergence of epistatic interactions, we address three key questions: (1) How does network complexity affect the maintenance of genetic variation under different recombination regimes? (2) What role do molecular components play in shaping adaptive trajectories? and (3) How do network-derived epistatic interactions influence the genetic architecture of adaptation? Our findings reveal unexpected relationships between network structure, genetic variation, and adaptive potential, with implications for understanding the evolution of complex traits in natural populations.

## Materials and methods

### Experimental design

We conducted a computational experiment to identify the effects of genetic architecture on adaptation to a phenotypic optimum. We explored 150 combinations of three genetic architecture parameters: the genome-wide recombination rate (*r*), the number of contributing loci (*n*_*loci*_), and the mutational effect size variance (*τ*). We evaluated ten recombination rate levels (*r* ∈ [10^−10^, 10^−9^, …, 10^−1^]), five numbers of loci (*n*_*loci*_ ∈ [4, 16, 64, 256, 1024]), and three treatments for the mutational effect size variance (*τ* ∈ [0.0125, 0.125, 1.25]). Recombination rates are expressed as the probability of crossover between two adjacent loci.

The range of recombination rates allowed us to investigate scenarios where genomic regions have suppressed recombination, such as centromeres or chromosomal inversions (Coop and Przeworski 2007) at the low end (e.g., *r* = 10^−10^), and where genes are on different chromosomes at the high end (*r* = 10^−1^), as well as more realistic scenarios with intermediate recombination rates. The number of loci treatments were chosen to explore oligogenic architectures (4, 16) and highly polygenic architectures (256 - 1024). Powers of four were used to ensure both the K+ and K− models were balanced in the number of loci contributing to each molecular component. The mutational effect size variance values were chosen to approximate small, intermediate, and large effect distributions of new mutations. In other words, the smaller *τ* is, the smaller the average effect of a given mutation. More detail is provided in S1 Appendix.

Each genetic architecture combination was evaluated in three models: an additive model and two treatments of the NAR network motif. These models differed in how the genotype translated to phenotype, with the NAR motif introducing curvature to the genotype-phenotype map. The NAR model is explained in detail in O’Brien *et al*. (2024). In summary, the NAR motif consists of two genes, *X* and *Z. X* expression is given by a step function activated by an environmental cue. *X*’s product activates *Z* and *Z*’s product represses further *Z* expression. The phenotype in this model is the total amount of *Z* expression in a fixed time interval, which is given by solving the ordinary differential equation (ODE) describing the NAR network’s dynamics. The change in *Z* expression with respect to time *t* is given by:

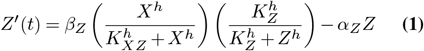

Here, *h* is the Hill coefficient (fixed at *h* = 8 for a steep activation curve after *X* is switched on; Alon 2019), and *α*_*Z*_, *β*_*Z*_, *K*_*Z*_, and *K*_*XZ*_ are molecular components: quantitative traits under direct control of QTLs. QTLs represent genes or regulatory regions of similar size and mutations at those loci represent sequence changes somewhere within that region. Variation in the molecular components drives variation in *Z* expression, and the resulting trait value and fitness. We solve for *Z* expression over time by expressing the ODE as an initial value problem and solving numerically with the Ascent header library (Berry *et al*. 2021). A step function gives *X* expression

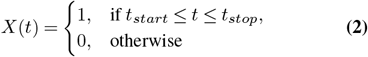

where *t*_*start*_ and *t*_*stop*_ are the boundaries where the *X*-activating environmental cue is present. We chose *t*_*start*_ = 1 and *t*_*stop*_ = 6 and solved for *Z* expression during the period *t* ∈ [0, 10], so *X* was active for half the total evaluated time period. The *Z* expression period was chosen to balance computational cost with a long enough time to capture a wide range of *Z* dynamics. For more detail on the parameterization of the ODE, see O’Brien *et al*. (2024).

The NAR treatments differed in which molecular components were mutable. In the K+ treatment, all four components (*α*_*Z*_, *β*_*Z*_, *K*_*Z*_, and *K*_*XZ*_) could evolve. In the K−treatment, *K*_*Z*_ and *K*_*XZ*_ were fixed at *K*_*Z*_ = *K*_*XZ*_ = 1. *K* values were chosen to match the *K* values used in O’Brien *et al*. (2024).

Fitness was given by Lande’s (1976) fitness function:

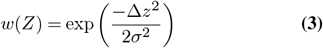

where Δ*z* is the distance to the phenotypic optimum, and *σ* is the width of the fitness function, representing selection strength. We set *σ* = 0.05, representing an approximately 5% drop in fitness following the optimum shift.

#### Fixed parameters

Among all treatments, some population parameters were kept constant. These included the mutation rate and population size. The per-locus, per-generation mutation rate, *µ* = 10^−5^, was chosen based on estimates of per-locus mutation rates from Lynch (2010), assuming 1000 sites per locus when converting from per-base-pair to per-locus mutation rates. *µ* represents the probability of a new mutation occurring at any given locus in a given gamete and generation. The mutation rate was adjusted for *n*_*loci*_ so that the genome-wide mutation rate was identical across genetic architectures (and so the effects of changing *n*_*loci*_ were related to the number of loci rather than changes in the mutation rate). The population size was fixed at 5,000 individuals to balance the effects of drift with simulation speed.

#### Simulations

Combined with the three models (Additive, K+, K−), our design produced 450 unique genetic architectures. We replicated each combination 48 times via randomly generated seeds. We input the genetic architecture parameter values into a forward-time Wright-Fisher simulation implemented in a custom version of SLiM 4.0.1 (Haller and Messer 2022). The SLiM version used is available at https://github.com/nobrien97/SLiM/releases/tag/PolygenicNAR2024. Our previous paper (O’Brien *et al*. 2024) provides more information on our implementation. Each replicate first adapted to an initial optimum phenotype for 50,000 generations of burn-in (*O*_*burn*−*in*_ = 1).

Then, the optimum was instantaneously shifted to a new optimum *O* = 2. We then tracked adaptation for 10,000 generations, sampling allelic frequencies and effect sizes, and phenotypic means and variances every 50 generations. We ran the simulations on the National Computational Infrastructure’s Gadi HPC system.

### Analysis

#### Pairwise epistasis and fitness effect estimation

We randomly sampled pairs of mutations and estimated fitness epistasis (*ϵ*_*w*_) between them. Fitness epistasis was calculated between two loci with derived alleles *A* and *B* using

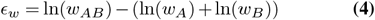

where *w*_*AB*_ is the fitness of the double mutant, and *w*_*A*_ and *w*_*B*_ are the fitnesses of the single mutants (individuals with only one of the two derived alleles). Fitness was measured on the ancestral background (i.e., no other segregating mutations), assuming that the wildtype has neither *A* nor *B*. Fitness was measured in diploid individuals and mutational effects measured in heterozygous state (i.e., AaBb individuals contributed to *w*_*AB*_, Aabb to *w*_*A*_ and aaBb to *w*_*B*_).

To estimate average fitness epistasis, we sampled 96 mutations with replacement from each simulation’s pool of segregating alleles, splitting the samples into two groups (A and B). We then calculated *ϵ*_*w*_ between A and B, for a total of 48 comparisons. We repeated this process 1,000 times for a total of 48,000 *ϵ*_*w*_ measures per simulation. We calculated the mean and standard deviation of *ϵ*_*w*_ across the 48,000 *ϵ*_*w*_ estimates and simulation replicates.

We also used this fitness data to assess the distribution of fitness effects. From the genotype data, we measured the proportion of beneficial mutations among mutations segregating in the population and the fitness effect size of those beneficial mutations.

The fitness effect *s*_*A*_ of mutation *A* was calculated as

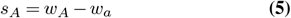

where *w*_*A*_ is the fitness of an individual with mutation *A*, and *w*_*a*_ is the fitness of individual lacking that mutation (but with an identical genotype across all other sites. As with epistasis, fitness was measured relative to the ancestral background (i.e. *w*_*a*_ is the wildtype genotype with no segregating mutations) and in heterozygous state.

To assess the effect of molecular component type on the proportion of beneficial mutations (*p*_*b*_), we used a beta regression model of form *p*_*b*_ ~ Molecular component, repeating this for K+ and K− models. We implemented the beta regression using the R package `betareg` 3.2.0 (Cribari-Neto and Zeileis 2010). For additive models, we computed the mean and 95% confidence intervals since additive models had only one molecular component.

We also measured the fitness effect size (*s*) of mutations in each model and molecular component. To measure the effects of molecular components, we used a generalized least squares (GLS) model of form s ~ Molecular component for K+ and K− models. This was implemented using the `gls()` function in package nlme 3.1.164 (Pinheiro and Bates 2006; Pinheiro *et al*. 2023). GLS was chosen to account for differences in variance between *s* estimates for each molecular component.

#### Pairwise linkage disequilibrium

To estimate pairwise LD between loci, we extracted the frequencies of minor alleles at all QTLs and the shared frequencies of those alleles across pairs of loci for each simulation. We removed all QTLs with minor allele frequency less than 5% and all multiallelic loci. This gave us up to 523,776 total pairwise comparisons in our largest case (when *n*_*loci*_ = 1024). We then calculated the genotype fitnesses of individuals with/without the minor alleles as per the methods given in the fitness epistasis section above (Eq. 4). We considered four genotypes, AB, aB, Ab, and ab, for each pair of QTLs, measuring fitnesses on a heterozygous background. The genotype AB had the highest fitness, ab had the lowest fitness, and the intermediates were Ab/aB. We used signed metrics to identify if the sign of epistasis was predictive of the sign of LD. We first used *D*′ but found that low-frequency alleles skewed the results towards extreme *D*′ values (−1 and 1). Instead, we calculated *D* (Lewontin 1964):

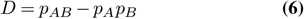

where *p*_*AB*_ is the frequency of the haplotype with the greatest fitness, and *p*_*A*_ and *p*_*B*_ are the marginal frequencies of the two minor alleles at loci *A* and *B*. As well as the overall pattern of LD, we calculated *D* between pairs of loci that shared similar minor allele frequencies (frequencies within 10% of each other). This gave an estimate of the pattern of LD across the site frequency spectrum.

#### Variance component estimation

We were also interested in the pattern of additive genetic variation under different genetic architectures and how variance was compartmentalized within the molecular components. To estimate variance components, we implemented a nested full-sib, half-sib crossing design (Lynch and Walsh 1998, pg 570 - 573). This design allows for accurate partitioning of the phenotypic variance into additive genetic, nonadditive genetic, and environmental components (Lynch and Walsh 1998). We randomly sampled five sires from the population at four timepoints during simulations (when the population first reached 25%, 50%, 75% and 100% of the required phenotypic change from burn-in to the new optimum). For each sire, we randomly sampled 20 dams and crossed each sire-dam combination 10 times to produce a total of *n*_ind_ = 1, 000 F1 individuals. Offspring sharing a sire and dam were full-sibs, and those sharing a sire were half-sibs. The F1 were genotyped and phenotyped (including measuring their molecular component values). In addition, the Wright relationship matrix of the F1 population was measured and stored. The F1 was destroyed at the end of this crossing experiment and did not cross with the evolving population to ensure the crossing experiment did not interfere with selection.

We used a multivariate linear mixed model to estimate the additive genetic variance/covariance between molecular components in the F1 population. We chose a Gaussian kernel regression approach by Xavier and Habier (2022) to reduce the complexity of regression coefficient estimation. The model took the form

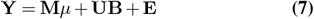

where **Y** is the *n*_ind_ *×n*_*C*_ phenotype matrix, *n*_ind_ is the number of individuals being predicted, *n*_*C*_ is the number of molecular components (either two for K− models or four for K+ models); **M***µ* is the *n*_ind_ *×n*_*C*_ matrix of intercept values; **U** is the *n*_ind_ *× n*_ind_ matrix of eigenvectors of the additive genetic relatedness matrix, **A**; **B** is the *n*_ind_ *× n*_*C*_ matrix of breeding values to be estimated; and **E** is the *n*_ind_ *×n*_C_ matrix of residuals.

Residuals were assumed to belong to a multivariate normal distribution **E** ~ 𝒩(**0, V**_**E**_), where *V*_*E*_ is the *n*_*C*_ *× n*_*C*_ diagonal matrix of residual variances for each molecular component. The model was implemented and solved in R 4.0.0 using the package `bWGR` 2.2.5 (Xavier and Habier 2022).

We also fit a second model to estimate *V*_*A*_ in *Z*. The same method was applied, however variance was calculated using variability in *Z* in **Y** instead of in the molecular components. Traits were scaled and centered before estimating the **G**_*C*_ matrix, however *Z* was not scaled/centered when calculating *V*_*A*_ in *Z* in the second model.

We extracted the estimated **G**_*C*_ matrix of additive genetic variances/covariances among the molecular components from the model. The regression was solved using a maximum likelihood estimator instead of the more conventional Bayesian MCMC techniques (Hadfield 2010). This was necessary owing to the large number of matrices to estimate, but this choice does increase the potential for error and reduces the ability to propagate error across analyses. Given the large number of matrices we estimated among our treatments and replicates, this error should be minimized when comparing means, however individual comparisons would be less reliable.

Because of the inherent error in estimating variance-covariance matrices, some estimates produced non-positive-semi-definite matrices. We used the method described by Higham (1988) to estimate the nearest positive definite matrix in these cases using the R package `Matrix` 1.7.0 (Bates *et al*. 2024).

#### G_C_ matrix analysis

To compare the additive variance and covariances among molecular components within and between the K+ and K− models, we used a variety of approaches characterizing the similarity between the size and shape of **G**_*C*_ matrices. We first used the metrics outlined in Hansen and Houle (2008), including evolvability, conditional evolvability, respondability, and autonomy. Evolvability measures the unconstrained response to selection in the direction of the selective pressure. This is predicted by the amount of genetic variance in the direction of selection (Hansen and Houle 2008). Conditional evolvability measures the response to directional selection in one molecular component, assuming that the other components are under stabilizing selection. It indicates the constraints on genetic variance within the **G**_*C*_ matrix.

Respondability is a measure of the total response to selection (including change outside the direction of selection), and autonomy is the proportion of additive variance in a molecular component that is independent of the other molecular components (Hansen and Houle 2008). Together, these metrics measure the ability of populations to respond to directional selection given the variation available to them among all molecular components. We computed these metrics via code/materials adapted from Hansen and Houle (2008) and Puentes *et al*. (2016). To identify the marginal effects of recombination and model type on the evolvability metrics, we used generalized least squares (GLS) models of the form response ~ model × recombination rate, where the recombination rate is one of *r* = 10^−10^, *r* = 10^−5^, or *r* = 10^−1^. We chose a GLS to account for differences in variance between model groups (K+ and K−). This was implemented using the `gls()` function in package `nlme` 3.1.164 (Pinheiro and Bates 2006; Pinheiro *et al*. 2023). To assess the significance of the predictors on the evolvability metrics, we conducted a type I ANOVA on each GLS fit using the `stats` 4.4.1 package (R Core Team 2023). We then calculated contrasts of the GLS fits using the R package `emmeans` 1.10.1 (Lenth 2023).

To compare the shape of the **G**_*C*_ matrices (i.e., the direction of variation among molecular traits), we first used complete linkage hierarchical clustering to compare Power-Euclidean distances (with power = 0.5) between **G**_*C*_ matrices:

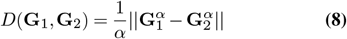

where || *X*|| is the Euclidean norm, **G**_1_ and **G**_2_ are covariance matrices and *α* is the power coefficient (Dryden *et al*. 2009). Power-Euclidean distance was chosen instead of Euclidean distance as it better matches the non-Euclidean space of **G**_*C*_ matrices, improving the accuracy of distance measures (Dryden *et al*. 2009). Power-Euclidean distance was calculated using the `distcov()` function in the R package `shapes` 1.2.7 (Dryden 2023). We constructed a tree using R packages `ape` 5.8, `tidytree` 0.4.6, and `phytools` 2.3.0 in R 4.4.1 (Paradis and Schliep 2019; Yu 2022; Revell 2024). This clustered **G**_*C*_ matrices with similar variance-covariance structure.

To corroborate the Power-Euclidean tree, we also calculated the PCA similarity factor between pairs of **G**_*C*_ matrices (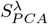; Singhal and Seborg (2005)). 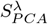 measures the similarity between principal components (PC) of pairs of **G**_*C*_ matrices, weighting differences in the angle between PCs by the size of the PCs (Yang and Shahabi 2004; Singhal and Seborg 2005). We used a bootstrap approach to repeatedly sample pairs of **G**_*C*_ matrices, noting the model (K+ or K−) and recombination rate for each matrix. For each matrix pair, we calculated 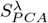 between the two via the equation:

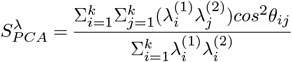

where *k* is the number of PCs (*k* = 4), *θ*_*ij*_ is the angle between the *i*th PC of the first **G**_*C*_ matrix and the *j*th PC of the second **G**_*C*_ matrix, and 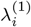 and 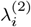 are the *i*th eigenvalues of the two **G**_*C*_ matrices (Singhal and Seborg 2005). We used the `PCASim()` function in R package `evolqg` 0.3.5 to calculate 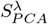 (Melo *et al*. 2015). We repeated this process for 10,000 bootstrap comparisons per model and recombination rate treatment. To analyze differences in PCA similarity between pairs of model types, we conducted a fractional logit regression with formula PCA_sim_ ~ model × recombination rate. We then assessed the significance of the model with a Chi-squared test implemented via in the `joint_tests()` function in `emmeans` 1.10.1 (Lenth 2023).

## Results

### Adaptability is a product of recombination, polygenicity, and effect size variability

We first explored differences in the adaptive response between different genetic architectures. We defined “adapted” populations as populations that reached within 5% of the new fitness optimum by the end of the simulation (i.e., adapted populations had mean phenotype 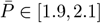). We found that under intermediate and large effect size variance regimes (*τ* ≥ 0.125), *>* 99% of replicates in all three models adapted within the 10,000 generation test period, regardless of the number of loci and recombination rate (Fig. S1, S2, S3). Under these effect sizes, increasing recombination reduced the adaptive rate of both NAR models and improved that of additive populations. However, this difference was minor except for under the *r* = 10^−1^ case (Fig. S4).

The largest differences between models were when populations were limited to mutations of small effect (*τ* = 0.0125). In this case, the rate of adaptation in additive and K−populations was slowed under low recombination (Fig 1). In addition, additive populations rarely reached the optimum: when *r* ≤ 10^−7^, only 8.22% of populations (n = 79) were able to adapt, whilst 98.6% of the K−populations (n = 947) reached the optimum under the same recombination treatment. As recombination increased, K− and additive models recovered a rapid adaptive rate. The response to recombination depended on the number of loci: when there were few QTL, increasing the recombination rate had little effect on the adaptation rate for all three models (Fig. 1). However, these findings were not shared with the K+ model. K+ populations were robust to low recombination rates, regardless of the number of loci (Fig. 1). We focus only on the *τ* = 0.0125 case for the remainder of the results to investigate this strong distinction between the K+ model and the other treatments. Similarly, because of the incremental differences seen with increasing recombination rate, we chose to compare models with *r* ∈ (10^−10^, 10^−5^, 10^−1^) for a low, medium, and high recombination rate treatment. In Table 1, we summarize our findings for these treatments.

**Table 1.** Summary of main results.

| Model | Recombination rate | Behavior |
| --- | --- | --- |
| K+ | High | Adaptation is rapid and linkage disequilibrium (LD) is low. Epistasis is positive and similar combinations of molecular components contribute to variation across replicates. |
| K+ | Low | Adaptation remains rapid, but LD is more extreme. Epistasis is positive and different combinations of molecular components contribute to variation across replicates. |
| K- | High | Adaptation is rapid (although slightly slower than K+), and LD is low. Epistasis is positive and similar combinations of molecular components contribute to variation across replicates. |
| K- | Low | Adaptation is slowed and LD is increased, although to a lesser extent than K+ models. Epistasis is positive and similar combinations of molecular components contribute to variation across replicates. |
| Additive | High | Adaptation is rapid and similar to K- models. LD is low. Epistasis is close to 0 (slightly positive). |
| Additive | Low | Adaptation is slow and rare – most populations do not reach the optimum. Populations so rarely adapt that we do not have enough samples for good estimates of LD. Epistasis remains close to 0. |

**Figure 1.**
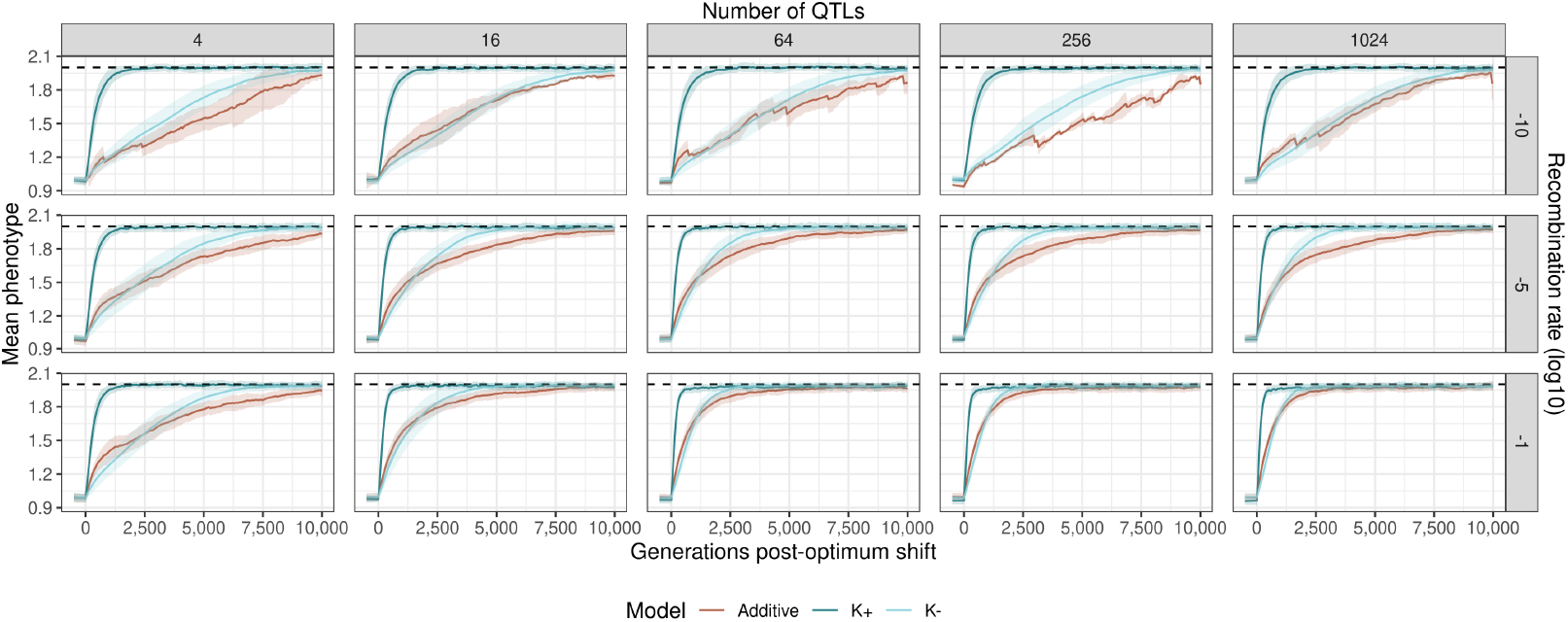
Adaptation to a shifting optimum in populations under small mutational effect size variance (*τ* = 0.0125). Each line is the mean of population means among populations that reached the optimum within 10,000 generations. There were up to 48 replicates per group. Ribbons show 95% confidence intervals. The dashed line shows the phenotypic optimum at *O* = 2.

### Additive variance is increased by recombination

The maintenance of the rapid adaptive response in K+ populations under low recombination and the recovery of the adaptive response in the other two models suggests a differential level of additive variance between the treatments. Fisher’s fundamental theorem states that the response to selection is proportional to the amount of additive variance in fitness that populations harbor (Fisher 1930), suggesting that K+ models should have high levels of *V*_*A*_ in *Z*. Similarly, *V*_*A*_ should increase in the other two models as recombination increases. Additive variance (*V*_*A*_) was maintained even under low recombination in the K+ model; this was not seen in the additive and K− models (Fig. 2).

**Figure 2.**
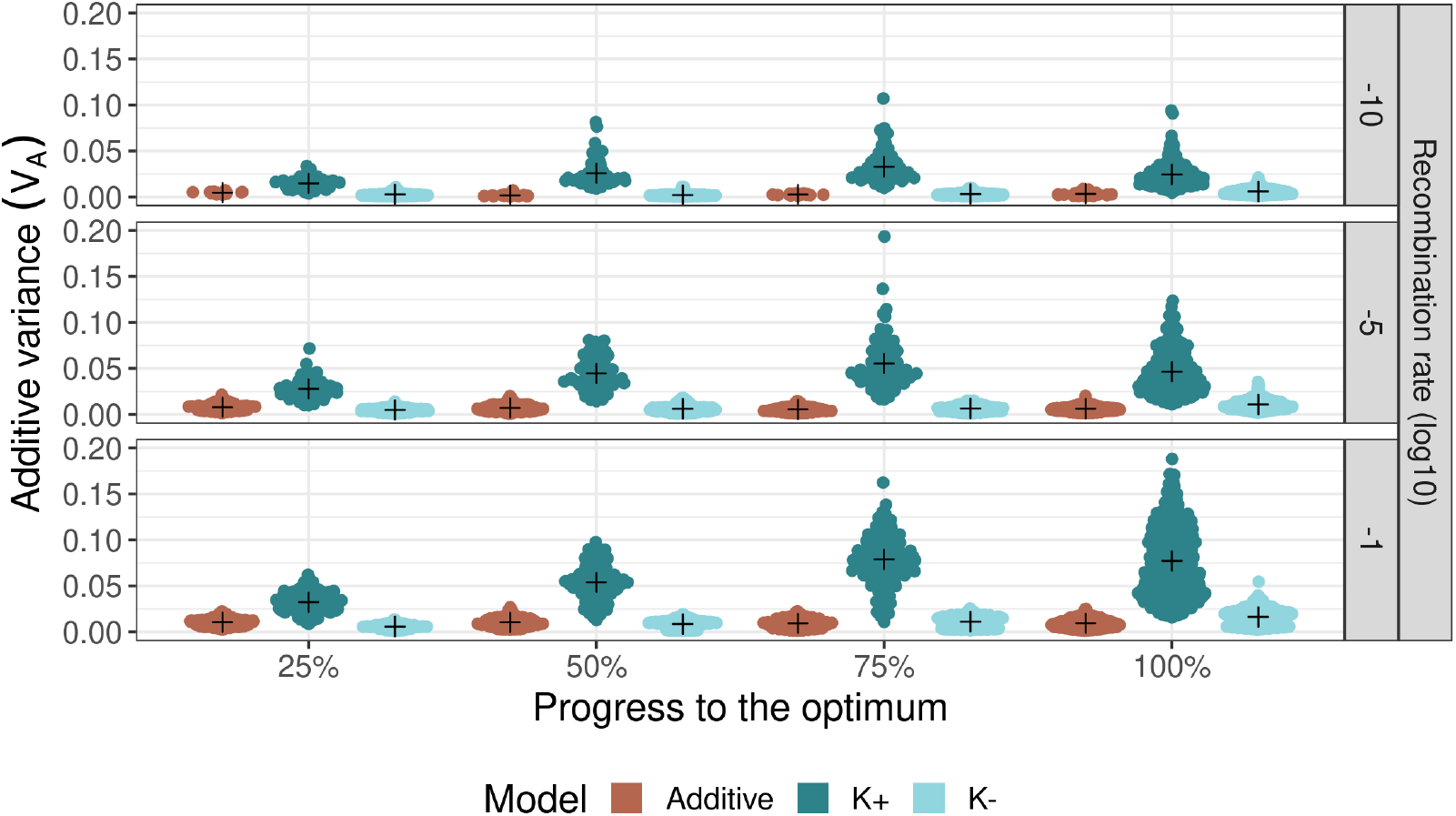
Additive variance (*V*_*A*_) in *Z* during adaptive walks in populations with small mutational effect size variance (*τ* = 0.0125). Each point represents a *V*_*A*_ estimate in a single population. The cross represents the mean *V*_*A*_.

The number of loci did not affect the amount of *V*_*A*_, and recombination’s effect on *V*_*A*_ was not contingent on the number of loci. In all models, the total change in *V*_*A*_ during the adaptive walk became less variable between replicates with increasing recombination, although the mean did not change.

In the additive and K+ models, the average change in *V*_*A*_ over the adaptive walk was around 0 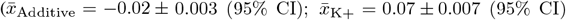, Fig S5), matching expectations for quantitative traits under selection (Barghi *et al*. 2020). However, *V*_*A*_ tended to increase over the adaptive walk in the K−model 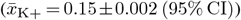. The increased additive variance in K+ populations and in high recombination rate treatments matched our expectations, but the origin of the difference between models was unclear. We considered that positive epistasis between strongly linked loci might have contributed to increased *V*_*A*_ in the K+ populations (Neiman and Linksvayer 2006). To investigate this, we evaluated the nature of pairwise epistatic interactions between segregating alleles in populations.

### Segregating alleles show synergistic epistasis in negative autoregulation simulations

We found that pairs of alleles showed positive fitness epistasis on average in all models, with K+ models harboring the largest amount (Fig. 3). Surprisingly, even the additive model showed slightly positive fitness epistasis 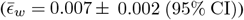, despite having zero trait epistasis and using a Gaussian fitness function with negative curvature that might be expected to generate negative epistasis in adapted populations. The epistasis signal was strongest in network models 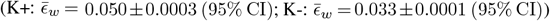, with differences between models maximized under low recombination. Given the positive nature of epistasis in NAR populations, which suggests that recombination would be deleterious (Otto and Whitlock 1997), we were interested in whether linkage disequilibrium matched the epistasis observed.

**Figure 3.**
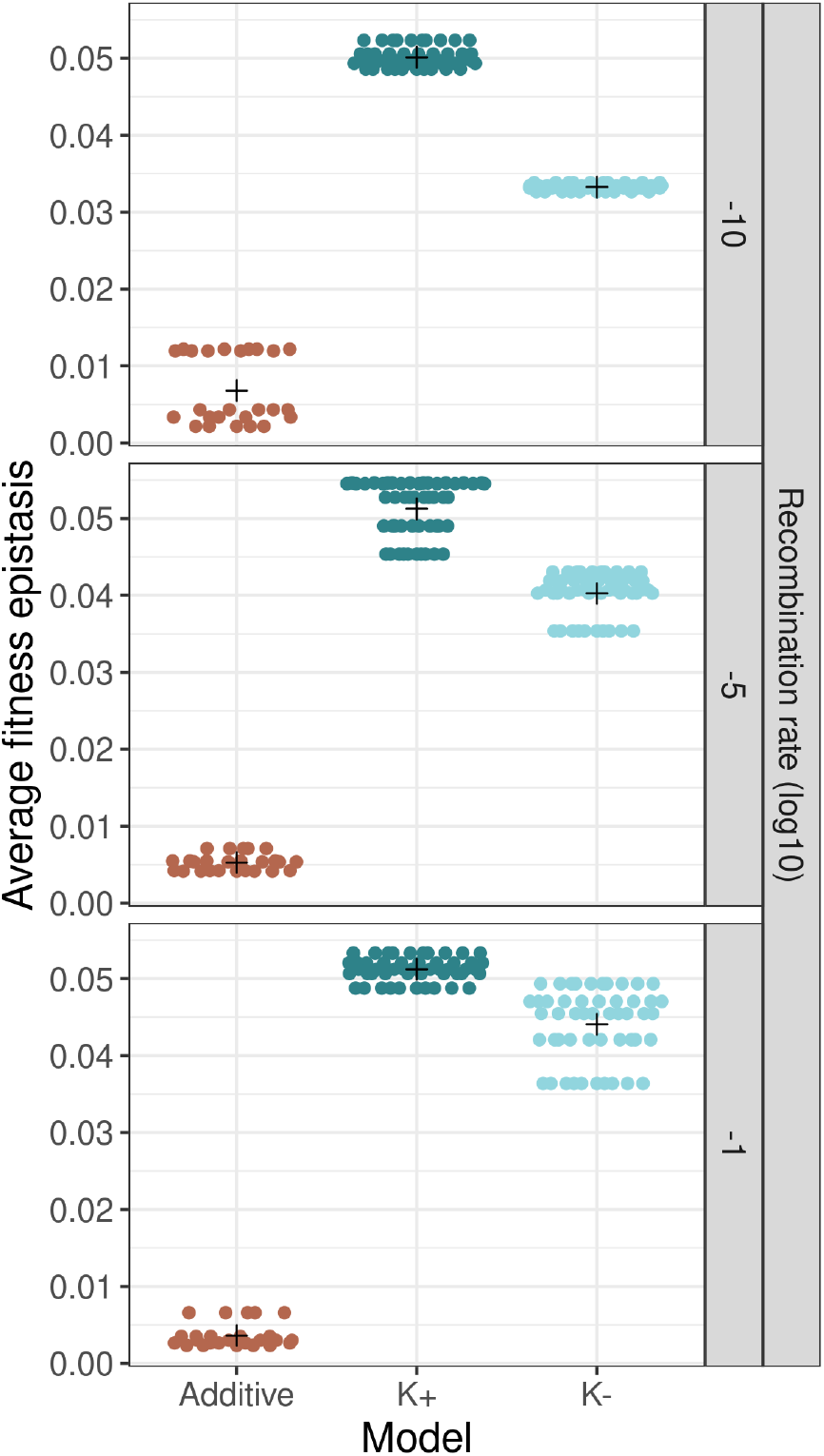
Mean pairwise fitness epistasis 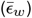 in adapted populations with small mutational effect size variance (*τ* = 0.0125). Each point represents 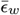 in a single population. The cross represents the mean 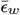 across different number of loci treatments and timepoints.

### Network complexity influences patterns of linkage disequilibrium

We found that pairwise LD was close to zero in all models under high recombination (Fig. 4A; 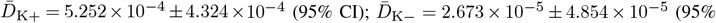 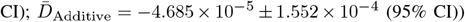, consistent with theoretical expectations. While positive epistasis was observed in our models, high recombination rates effectively broke down LD between loci. Since very few additive populations could adapt in low recombination combinations (under the small effect size treatment), we are not confident in concluding any differences in LD owing to recombination in additive models.

In NAR models, we found that the number of loci was uninformative in terms of the distribution of LD. Instead, differences in the distribution of LD were driven by model type and recombination rate. The K+ model appeared to harbor marginally more extreme LD (both positive and negative) than the K−model at low recombination (*r* = 10^−10^) (Fig. 4A), with the average variance in *D* across replicates being 0.007*±* 0.002 (95% CI) for K+ models and 0.004 ± 0.001 (95% CI) for K− models. This difference persisted as recombination increased (albeit to a lesser extent as recombination was still able to split extreme LD pairs: 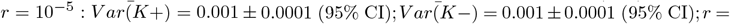 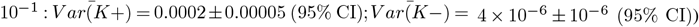. This trend persisted when we analyzed LD among pairs of QTLs with similar minor allele frequencies (Fig. 4B). Increasing recombination reduced the magnitude of LD across all models, as expected (Fig. 4).

**Figure 4.**
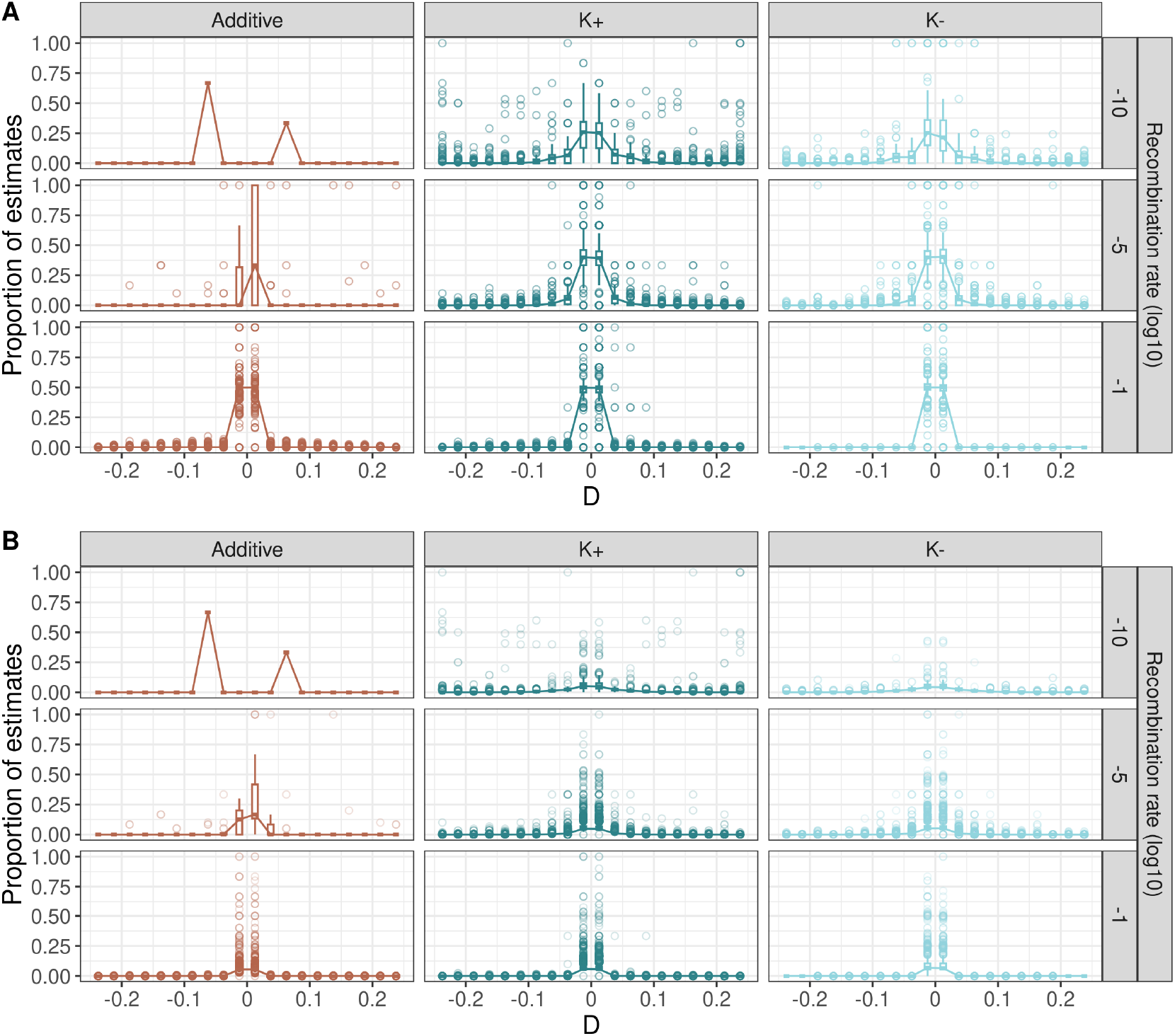
Distribution of linkage disequilibrium amongst pairs of QTLs. Boxes show variation across replicates and loci in pairwise Lewontin’s *D* estimates. *D* was split into 20 bins from −0.25 to 0.25 to plot boxes. Values are the proportion of *D* estimates in a given bin within a replicate. Due to bias concerns, QTLs with minor allele frequency *p* < 0.05 are not included. (A) shows the distribution of LD amongst all pairs of QTLs, regardless of their frequencies. (B) shows the distribution of LD amongst pairs of QTLs with minor allele frequency within 10% of each other to correct for estimation bias due to comparisons between rare and common QTLs.

Pairwise LD appeared to show combinations of alleles were more likely to be strongly associated in K+ models than additive and K− models. However, the direction of LD was not always aligned with epistasis. This disconnect could arise through Hill-Robertson effects, where genetic drift and interference between selected loci can mask the effect of beneficial alleles. When drift is strong relative to selection, beneficial alleles may be lost if they initially occur on disadvantageous genetic backgrounds, or deleterious alleles may hitchhike to high frequency when linked to beneficial mutations (Hill and Robertson 1966). Moreover, the increase in the proportion of QTLs under LD in K+ models might suggest that additional network complexity produces selection that maintains favorable allelic combinations. To identify the nature of these constraints, we considered whether the structure of additive variance-covariance in the networks’ molecular components was similar between K+ and K− models by comparing **G**_*C*_ matrices between populations.

### Additive variance-covariance in molecular components strongly differentiates K+ and K− models

We computed Power-Euclidean distances between all estimated **G**_*C*_ matrices to identify patterns in the variance structure of molecular components. Hierarchical clustering revealed strong clusters separating K+ and K− models (Fig. 5). While K+ and K− models fit into separate clusters, it was apparent that K− models harbored less variability overall (Fig. 5). This was confirmed with further analysis of the shape and size of **G**_*C*_ matrices (Fig. 6, Tables 2, 3, 4, 5) and PCA similarity (Fig. 7). We found strongly significant effects of the model/recombination rate combination on evolvability (*F*_2,6927_ = 67.5, *p* < 0.0001), conditional evolvability (*F*_2,6927_ = 248.5, *p* < 0.0001), respondability (*F*_2,6927_ = 405.4, *p* < 0.0001), and autonomy (*F*_2,6927_ = 1778, *p* < 0.0001). The number of loci had no effect on the evolvability metrics.

**Table 2.** Estimated mean evolvability from generalized least squares model fitting model type and recombination rate to evolvability. SE is the standard error, df is the degrees of freedom, and Lower and Upper CLs are the confidence limits. Means are calculated from the least squares model as marginal means.

| Model | Recombination rate | Mean evolvability | SE | df | Lower CL | Upper CL |
| --- | --- | --- | --- | --- | --- | --- |
| K+ | $10^{-10}$ | 0.9224 | 0.0018 | 942.20 | 0.9189 | 0.9259 |
| K- | $10^{-10}$ | 0.8885 | 0.0028 | 967.22 | 0.8831 | 0.8940 |
| K+ | $10^{-5}$ | 0.8797 | 0.0022 | 954.70 | 0.8754 | 0.8840 |
| K- | $10^{-5}$ | 0.8685 | 0.0031 | 1275.08 | 0.8625 | 0.8745 |
| K+ | $10^{-1}$ | 0.8405 | 0.0021 | 1161.54 | 0.8364 | 0.8446 |
| K- | $10^{-1}$ | 0.8629 | 0.0029 | 928.00 | 0.8573 | 0.8686 |

Degrees-of-freedom method: Satterthwaite
Confidence level used: 0.95

**Table 3.** Estimated mean conditional evolvability from generalized least squares model fitting model type and recombination rate to conditional evolvability. SE is the standard error, df is the degrees of freedom, and Lower and Upper CLs are the confidence limits. Means are calculated from the least squares model as marginal means.

| Model | Recombination rate | Mean conditional evolvability | SE | df | Lower CL | Upper CL |
| --- | --- | --- | --- | --- | --- | --- |
| K+ | $10^{-10}$ | 0.5514 | 0.0054 | 932.02 | 0.5407 | 0.5620 |
| K- | $10^{-10}$ | 0.7920 | 0.0048 | 1287.55 | 0.7827 | 0.8013 |
| K+ | $10^{-5}$ | 0.7500 | 0.0031 | 959.43 | 0.7438 | 0.7561 |
| K- | $10^{-5}$ | 0.8488 | 0.0033 | 1295.06 | 0.8423 | 0.8553 |
| K+ | $10^{-1}$ | 0.8002 | 0.0022 | 1155.12 | 0.7959 | 0.8045 |
| K- | $10^{-1}$ | 0.8608 | 0.0029 | 1335.99 | 0.8551 | 0.8665 |

Degrees-of-freedom method: Satterthwaite
Confidence level used: 0.95

**Table 4.**
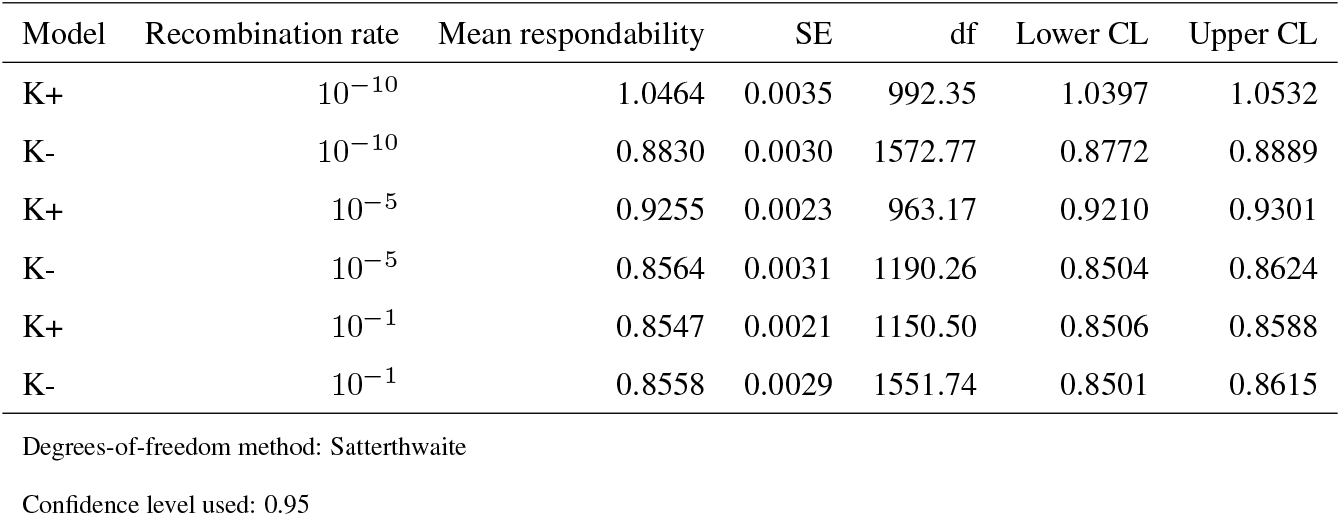
Estimated mean respondability from generalized least squares model fitting model type and recombination rate to respondability. SE is the standard error, df is the degrees of freedom, and Lower and Upper CLs are the confidence limits. Means are calculated from the least squares model as marginal means.

**Table 5.** Estimated mean autonomy from generalized least squares model fitting model type and recombination rate to autonomy. SE is the standard error, df is the degrees of freedom, and Lower and Upper CLs are the confidence limits. Means are calculated from the least squares model as marginal means.

| Model | Recombination rate | Mean autonomy | SE | df | Lower CL | Upper CL |
| --- | --- | --- | --- | --- | --- | --- |
| K+ | $10^{-10}$ | 0.6799 | 0.0054 | 956.76 | 0.6694 | 0.6904 |
| K- | $10^{-10}$ | 1.0052 | 0.0033 | 1241.85 | 0.9987 | 1.0117 |
| K+ | $10^{-5}$ | 0.8822 | 0.0021 | 959.28 | 0.8781 | 0.8862 |
| K- | $10^{-5}$ | 1.0344 | 0.0008 | 1293.68 | 1.0329 | 1.0359 |
| K+ | $10^{-1}$ | 0.9627 | 0.0005 | 1155.97 | 0.9617 | 0.9637 |
| K- | $10^{-1}$ | 1.0168 | 0.0004 | 1336.90 | 1.0160 | 1.0176 |

Degrees-of-freedom method: Satterthwaite
Confidence level used: 0.95

**Figure 5.**
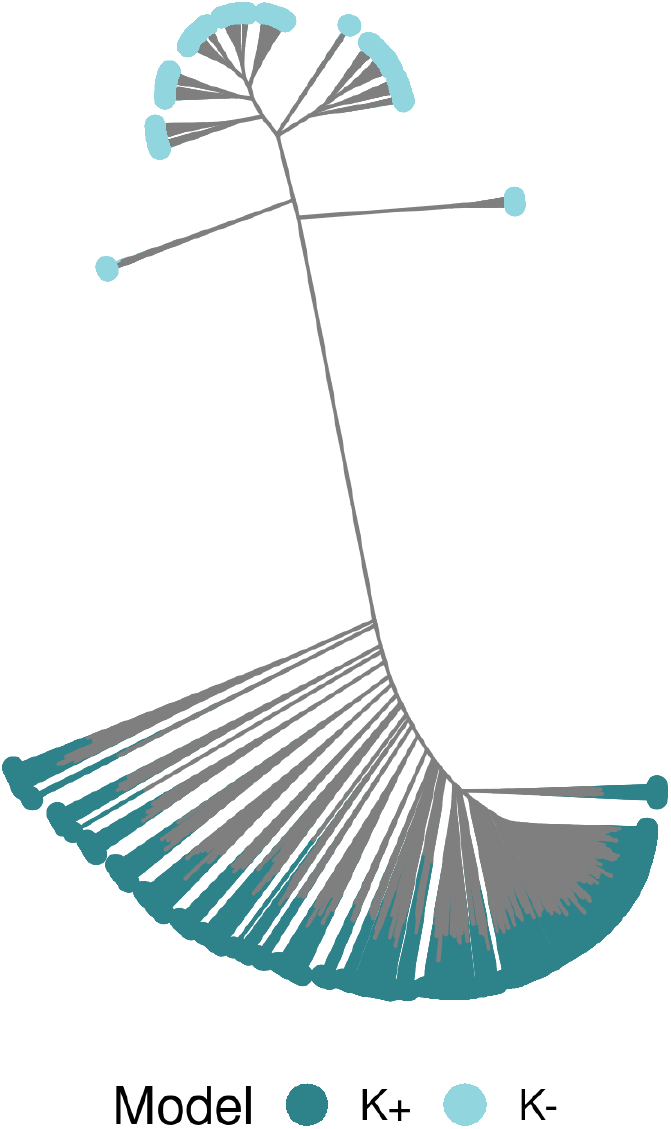
Distance tree of **G**_*C*_ matrices clustered by model type. Nodes represent populations and the distance between nodes represents their similarity. The tree is built on a hierarchical clustering model using Power-Euclidean distance (power = 0.5).

**Figure 6.**
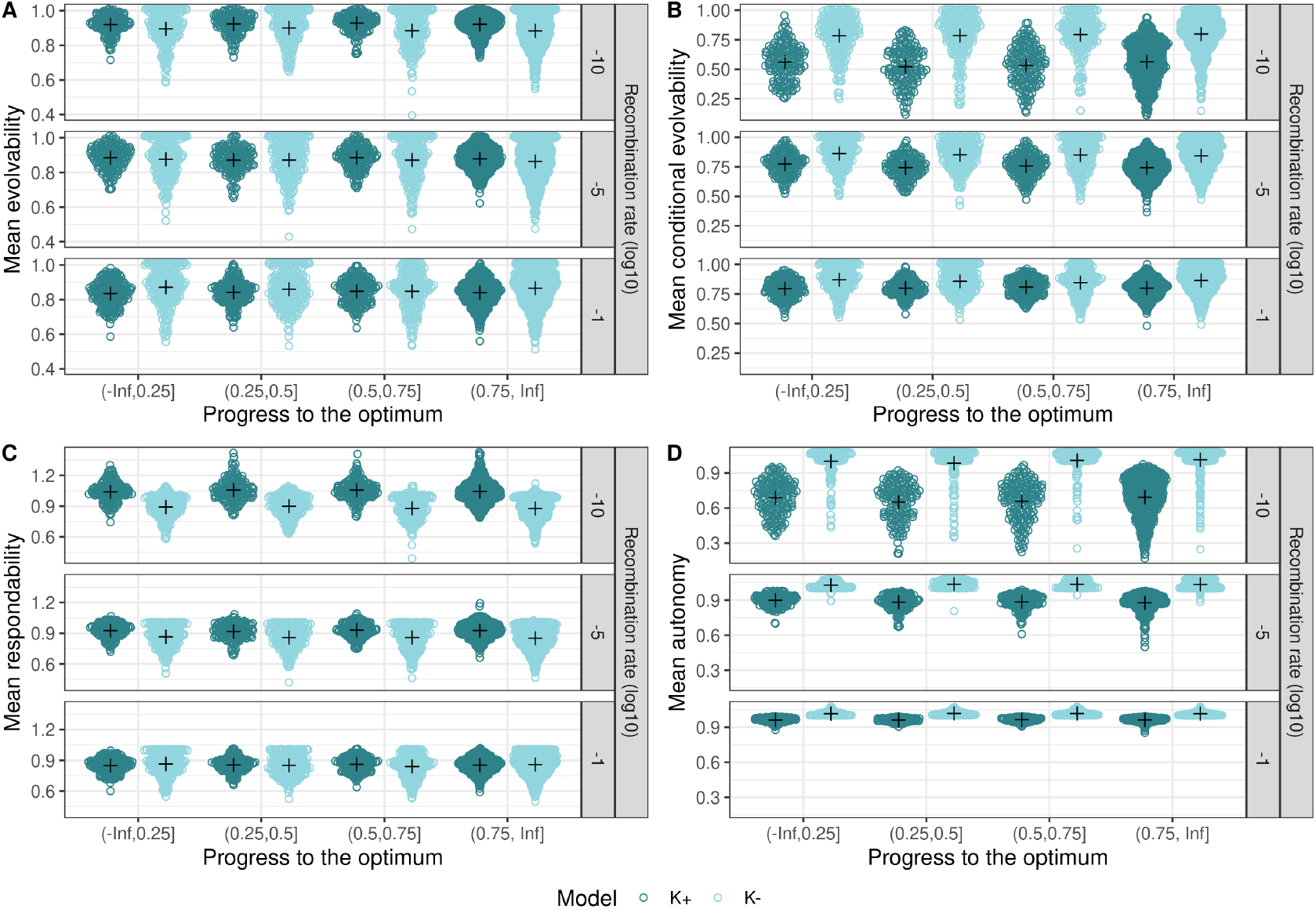
Evolvability metrics for **G**_*C*_ matrices in K+ and K− models. Each point represents a population and crosses represent group means. (A) shows mean evolvability, a measure of the total variation in molecular components. (B) shows the mean conditional evolvability, a measure of how much each molecular component can respond to directional selection if the other components are under stabilizing selection. (C) shows the mean respondability, the expected response to directional selection. (D) shows the mean autonomy, the average amount of independent variance across molecular components.

**Figure 7.**
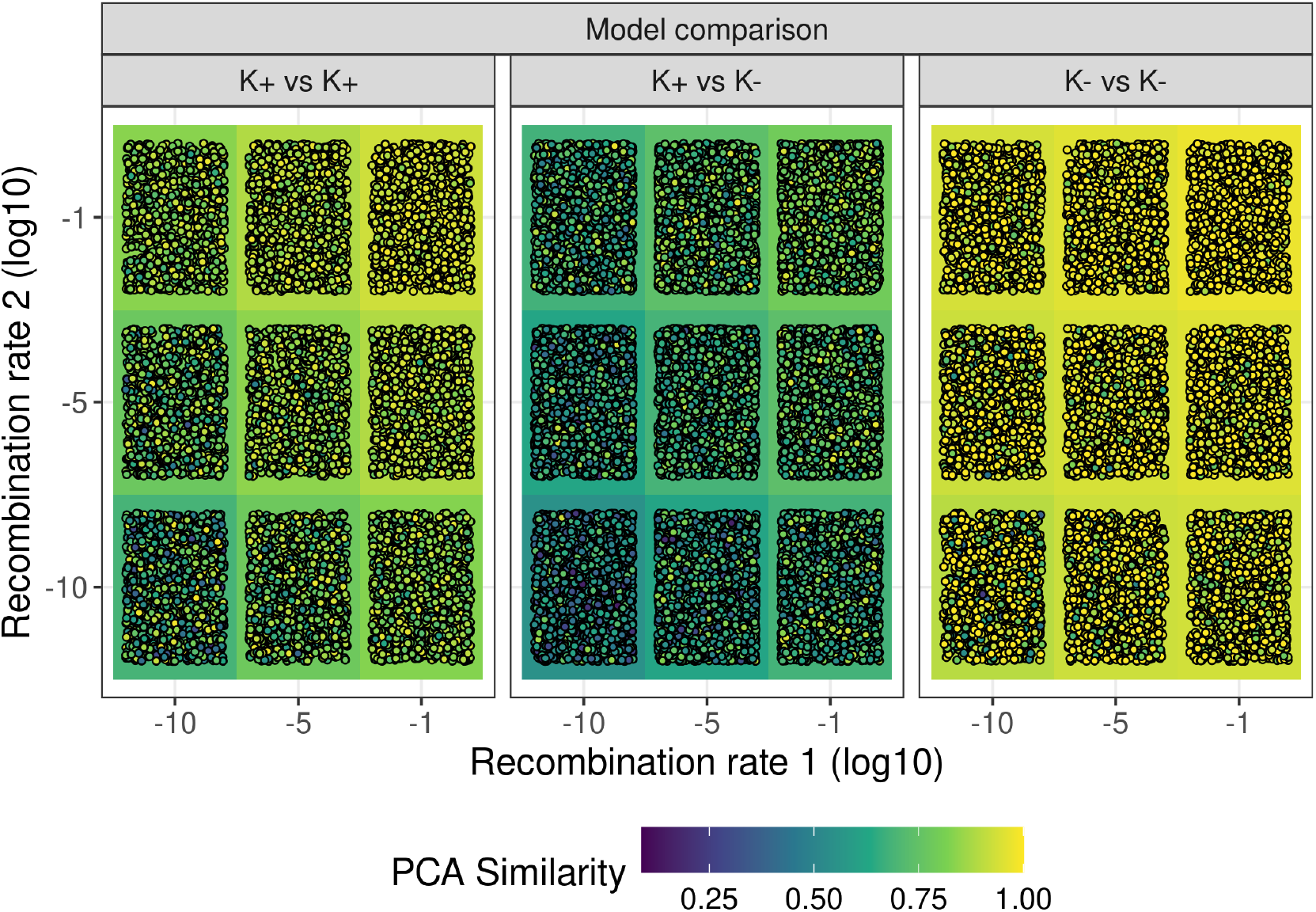
Principal components similarity between pairs of **G**_*C*_ matrices. Each point represents a bootstrap sample of two matrices. PCA similarity scores are correlations, with 0 meaning that matrices are completely independent with respect to genetic variation/covariation, and 1 meaning the **G**_*C*_ matrices are identical. Model combination refers to the model identity of the two matrices: both K+, both K−, or one of each. The x and y axes are the two recombination rates of the matrices. The background color is the average PCA similarity across all group samples.

Mean evolvability was roughly similar under all recombination rates (Table 2, Fig 6A). Under low recombination, K+ models had reduced mean conditional evolvability (Table 3, 6B), and mean autonomy (Table 5, 6D) compared to K− models, however, this effect diminished under higher recombination rates (Table 5, 6B,D). Mean respondability, however, was increased in K+ models under a low recombination rate (Table 4, 6C). Under high recombination, differences between the models in all evolvability metrics evaporate. This suggests that correlations between molecular components limit the amount of variation in any one molecular component in K+ models. The network prescribes a need for multiple components to be maintained simultaneously under low recombination.

**Table 6.**
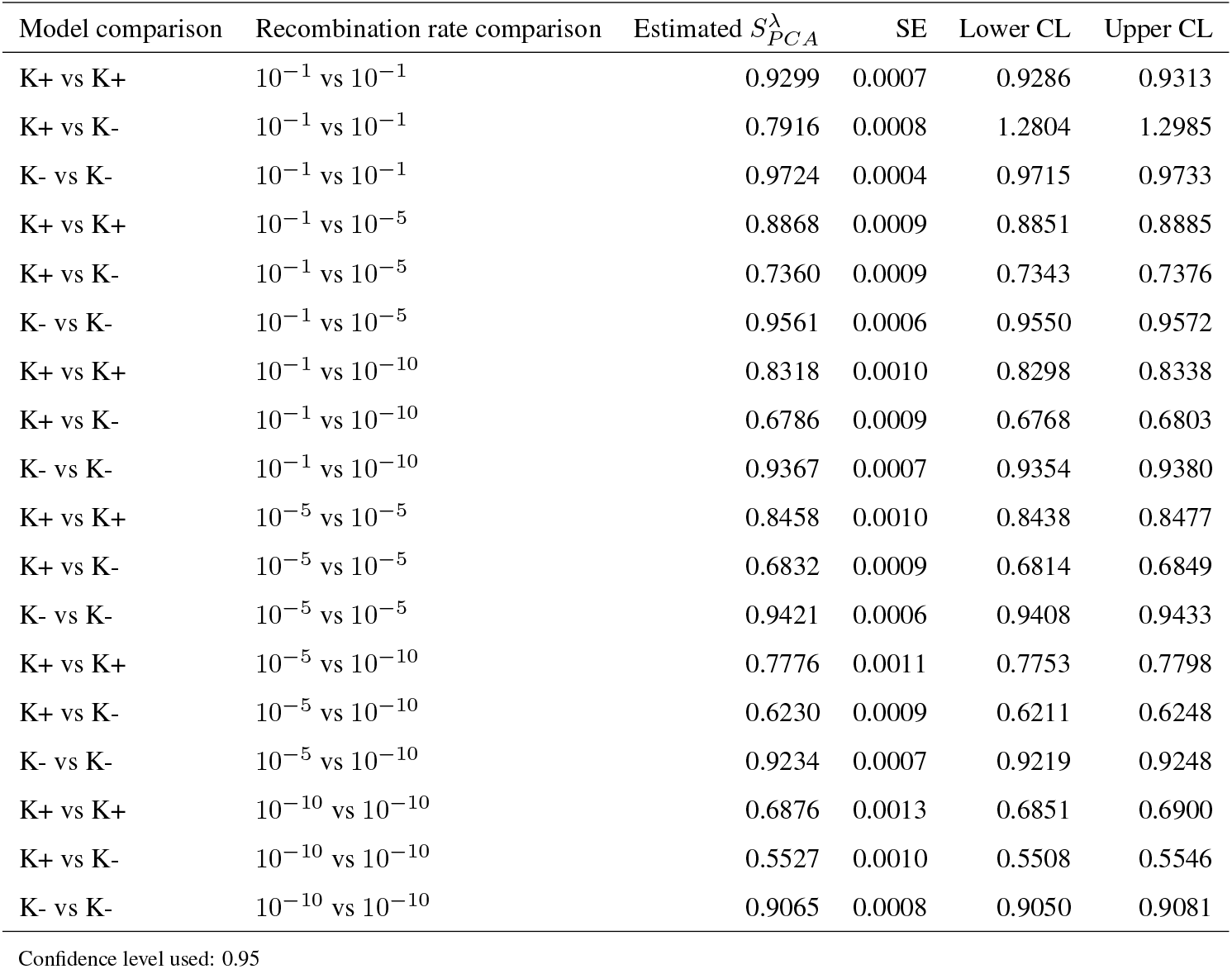
Estimated mean PCA similarity factor 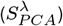 from a fractional logit regression of PCA similarity computed by pairs of **G**_*C*_ matrices differing in model composition and recombination rate. When 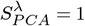, the two **G**_*C*_ matrices are identical, when 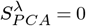, the two **G**_*C*_ matrices are orthogonal. SE is the standard error, and Lower and Upper CLs are the confidence limits. Confidence limits are asymptotic. Reciprocal comparisons have been removed from the table as they are not informative.

We analyzed the shape of variability in adapting populations further using the PCA similarity factor, 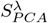, between randomly sampled **G**_*C*_ matrices (Singhal and Seborg 2005). This method calculates the similarity of direction and scale of the principal components of a covariance/correlation matrix. When 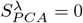, the matrices are orthogonal, sharing no variation. When 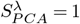, the matrices are identical (Yang and Shahabi 2004). We found that pairs of K−matrices were most similar across all recombination rates, comparisons of a K+ and a K−model were most dissimilar, and comparisons of two K+ matrices intermediate (Fig. 7; Model comparison:Recombination rate comparison interaction: 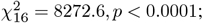, Table 6).

In all three comparisons, increasing recombination rates increased the similarity of the two matrices. This was exacerbated in comparisons with one or more K+ matrices, suggesting that the K+ models were more likely to use different combinations of molecular components (in different directions and different amounts of variance) to adapt, particularly when recombination was low. K− models were much more similar in their response, having fewer molecular components to organize. Together, the evolvability metrics and PCA similarity factor indicated that K+ populations tended to use different combinations of molecular components to adapt across replicates under low recombination.

To identify why this occurred, we investigated the distribution of fitness effects among mutations in each molecular component.

We studied two aspects of the distribution of fitness effects among molecular components: the proportion of beneficial mutations and the effect sizes of those beneficial mutations. Recombination did not affect the probability a segregating mutation was beneficial. Across all models, mutations were beneficial around 50% of the time (Fig 8, Table 7). This reflects the expectation under Fisher’s geometric model: the probability a mutation is beneficial approaches 50% as its effect approaches 0 (Fisher 1930). Since mutations were sampled from a narrow distribution (*τ* = 0.0125), and our fitness function was derived from the geometric model (Eq. 3, Lande 1976), meeting this expectation was unsurprising.

**Table 7.** Estimated mean proportion of beneficial mutations among all segregating mutations (*p*_*b*_). Estimates are marginal means from beta regressions of the proportion of beneficial mutations with different mutation types. Each model is a separate beta regression. SE is the standard error, and Lower and Upper CLs are the confidence limits. Confidence limits are asymptotic. The additive model is the mean among all mutations *±* the 95% confidence interval as there is only one mutation type and no need for a regression test.

| **Model: K-** |  |  |  |  |
| --- | --- | --- | --- | --- |
| Mutation Type | Estimated $p_b$ | SE | Lower CL | Upper CL |
| $\alpha_Z$ | 0.5279 | 0.0007 | 0.5265 | 0.5293 |
| $\beta_Z$ | 0.5295 | 0.0007 | 0.5281 | 0.5309 |
| **Model: K+** |  |  |  |  |
| $\alpha_Z$ | 0.5025 | 0.0013 | 0.5000 | 0.5050 |
| $\beta_Z$ | 0.5048 | 0.0013 | 0.5023 | 0.5073 |
| $K_Z$ | 0.0035 | 0.0001 | 0.0034 | 0.0037 |
| $K_{XZ}$ | 0.5358 | 0.0013 | 0.5334 | 0.5383 |
| **Model: Additive** |  |  |  |  |
| Mutation Type | Mean | SE | Lower CL | Upper CL |
| Additive | 0.4928 | 0.0016 | 0.4898 | 0.4959 |

Confidence level used: 0.95

**Figure 8.**
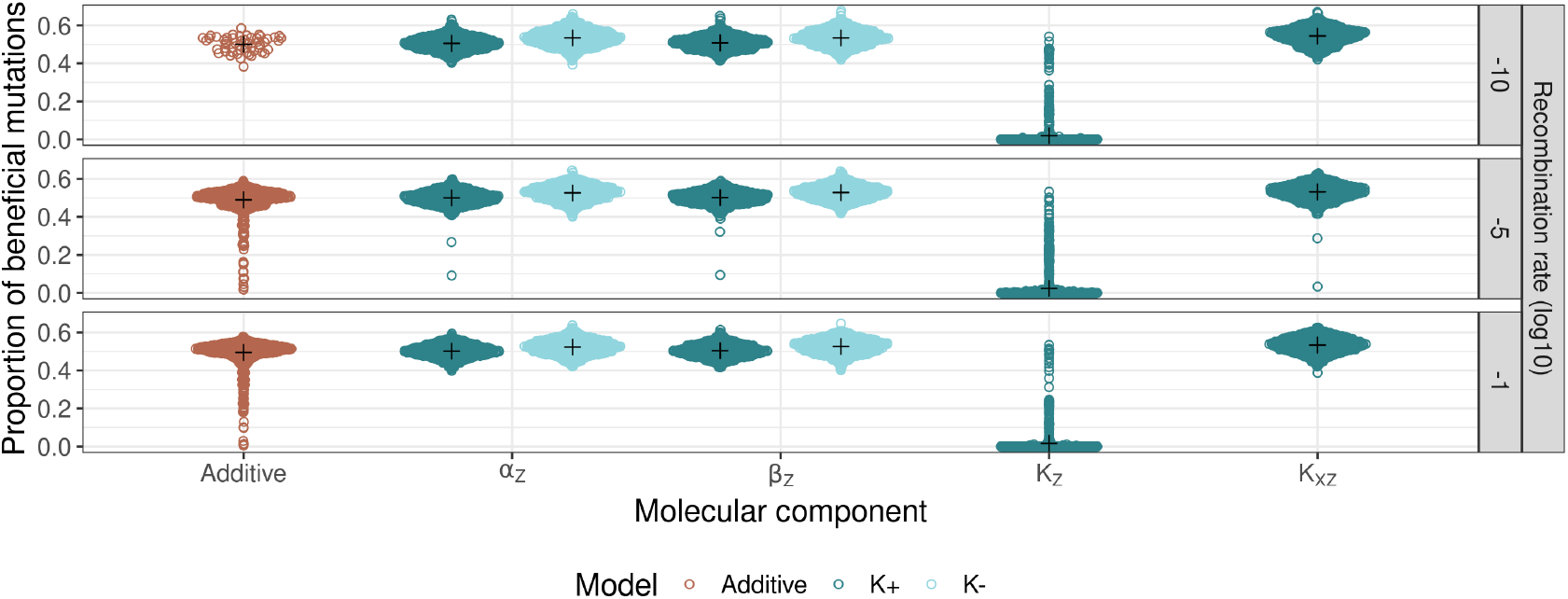
Proportion of mutations that are beneficial per model/molecular component and recombination rate. Each point represents a mutation and crosses represent group means. Beneficial mutations were sampled from populations during the walk, and fitness was calculated depending on their genetic background.

The proportion of beneficial mutations did not differ between molecular components in K− models (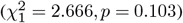. However, it did in K+ models (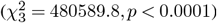,. In K+ models, mutations affecting *K*_*Z*_ were extremely unlikely to be beneficial (Fig 8, Table 7). We additionally examined the fitness effects of beneficial mutations, finding they differed between molecular components in the K+ model (*F*_1,1287754_ = 738220.8, *p* < 0.0001), but not in the K−model (*F*_1,857718_ = 0.1, *p* = 0.714). In the K+ model, beneficial *K*_*Z*_ mutations had were close to neutral. In contrast, *K*_*XZ*_ mutations were more strongly beneficial on average, but with greater variability in their effect (Fig. 9, Table 8). This suggests that adaptation in the K+ model is a combination of small effect *α*_*Z*_ and *β*_*Z*_ mutations and larger effect *K*_*XZ*_ mutations.

**Table 8.** Estimated mean fitness effects among beneficial mutations across different mutation types and models. In K+ and K− models, estimates are the estimated marginal means of fitness effects from generalized least squares models of fitness effects across different mutation types and models. Each model is a separate test. K+ models differed in effect sizes between mutation types, whereas K− models did not. K− and additive tables give means and 95% confidence limits, whereas the K+ model gives estimated marginal means and asymptotic 95% confidence limits. SE is the standard error, df is the degrees of freedom, and Lower and Upper CLs are confidence limits. Confidence limits are asymptotic. The additive model estimate is the mean among all mutations *±* the 95% confidence interval as there is only one mutation type.

| **Model: K-** |  |  |  |  |  |
| --- | --- | --- | --- | --- | --- |
| Mutation Type | Mean fitness effect | SE | df | Lower CL | Upper CL |
| $\alpha_Z$ and $\beta_Z$ | 0.00188 | 0.000002 | NA | 0.001877 | 0.001883 |
| **Model: K+** |  |  |  |  |  |
| Mutation Type | Estimated marginal mean | SE | df | Lower CL | Upper CL |
| $\alpha_Z$ | 0.0018 | <0.0001 | 1287751 | 0.0018 | 0.0018 |
| $\beta_Z$ | 0.0018 | <0.0001 | 1287751 | 0.0018 | 0.0018 |
| $K_Z$ | 0.000001 | <0.0001 | 1287751 | <0.00001 | <0.00001 |
| $K_{XZ}$ | 0.0055 | <0.0001 | 1287751 | 0.0055 | 0.0055 |
| **Model: Additive** |  |  |  |  |  |
| Mutation Type | Mean | SE | df | Lower CL | Upper CL |
| Additive | 0.000443 | 0.000001 | NA | 0.000442 | 0.000444 |

Confidence level used: 0.95

**Figure 9.**
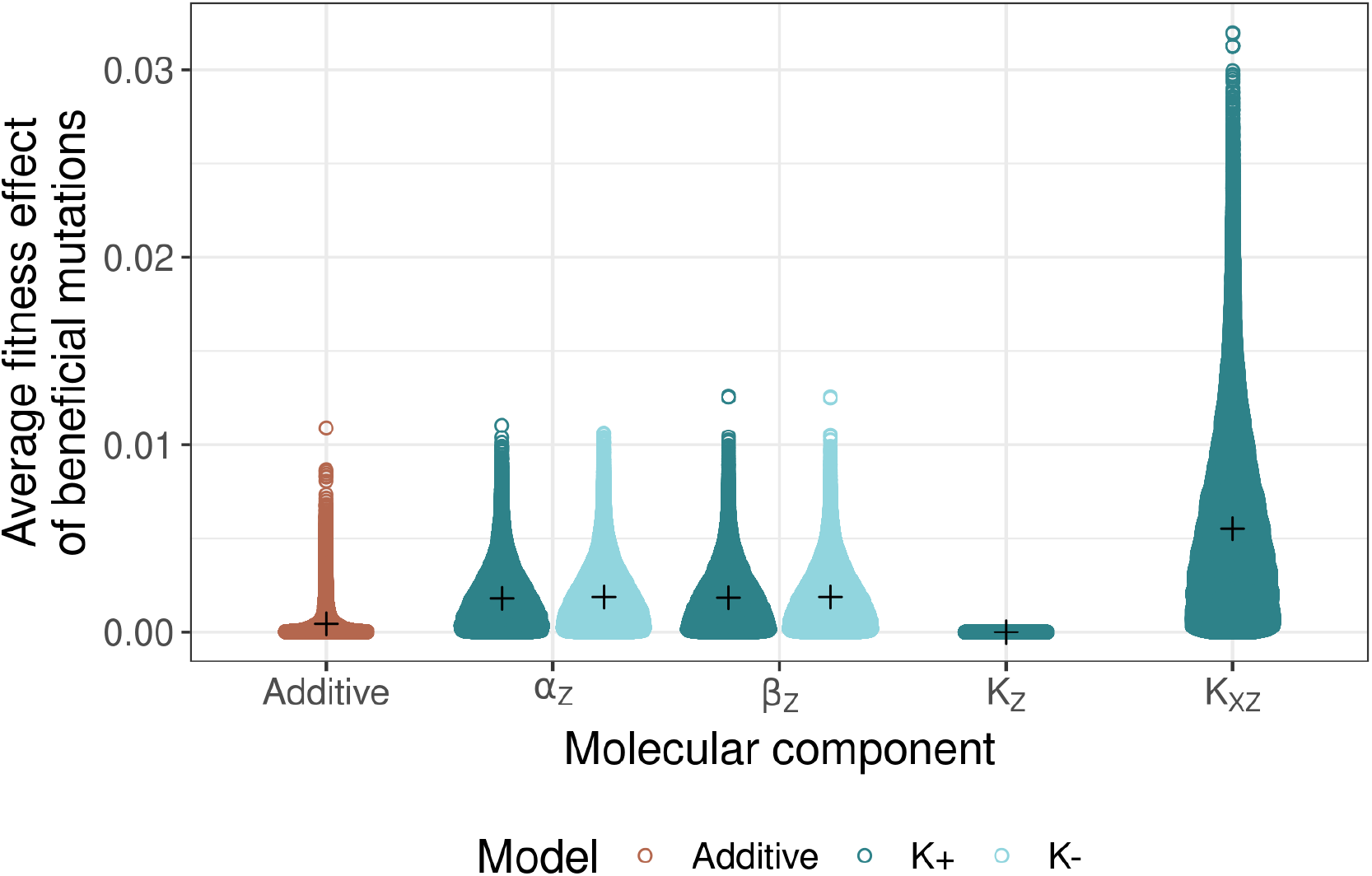
Average fitness effect of beneficial mutations across models/molecular components. Each point represents a mutation, crosses represent group means.

## Discussion

Our study demonstrates that negative autoregulation network architecture fundamentally influences both adaptive trajectories and the maintenance of additive genetic variance. These effects are modulated by three key parameters: recombination rate, mutational effect size, and the number of contributing loci (Fig. 1, Fig. S1, S2, S3). The molecular architecture of traits, particularly through the partitioning of additive variance, constrains the exploration of phenotypic space (Fig. 7, 9), with this constraint most pronounced when genetic interactions promote specific allelic combinations.

Our most striking finding emerged in highly polygenic architectures with small mutational effects: K+ populations maintained rapid adaptive responses under low recombination, while K− and additive populations exhibited impaired adaptation (Fig. 1). This adaptive advantage coincided with elevated additive variance in *Z* expression (Fig. 2) and distinct patterns of variance partitioning among molecular components. Below, we discuss the biological mechanisms driving the robustness to low recombination in K+ models, how our findings interface with existing models of evolution and with empirical results, and how gene regulatory networks might contribute to the evolution of recombination.

### Network Structure and Adaptation

The differential adaptive responses between K+ and K- models stem from their distinct architectures of additive genetic variance within the *Z* expression network. In K+ models, two additional parameters (*K*_*Z*_ and *K*_*XZ*_) contribute to phenotypic variation beyond the basic *α*_*Z*_ and *β*_*Z*_ components. Our analysis revealed that *K*_*XZ*_ mutations generated significantly larger fitness effects compared to other molecular components, including *α*_*Z*_ and *β*_*Z*_ (Fig. 9). In contrast, *K*_*Z*_ mutations were predominantly neutral, suggesting they did not contribute substantially to the increased epistasis and variable linkage disequilibrium observed in K+ populations (Fig. 3, 4). This asymmetry in mutational effects between *K*_*Z*_ and *K*_*XZ*_ emerges directly from the mathematical structure of the underlying ordinary differential equation.

In Eq. 1, *K*_*Z*_ represents the concentration of *Z* product required to reach half the maximum autoregulation strength (Alon 2019). Smaller *K*_*Z*_ values result in stronger autoregulation and vice versa. Given our optimum choice, to change the amount of autoregulation meaningfully requires large effect *K*_*Z*_ mutations. Such mutations are unlikely to be sampled in the *τ* = 0.0125 treatment (Fig. 10A). Hence, most *K*_*Z*_ mutations we observed were neutral. This behavior matches empirical work by Kozuch *et al*. (2020), who found the steady-state concentration of LexA in *E. coli* was sensitive to *K*_*Z*_ only when (*β*_*Z*_*/α*_*Z*_)*/K*_*Z*_ *>* 1. Even in this case, the effect of *K*_*Z*_ on expression required small values of *K*_*Z*_ (*K*_*Z*_ *<* 0.6) (Kozuch *et al*. 2020), well outside the *τ* = 0.0125 case we explored.

**Figure 10.**
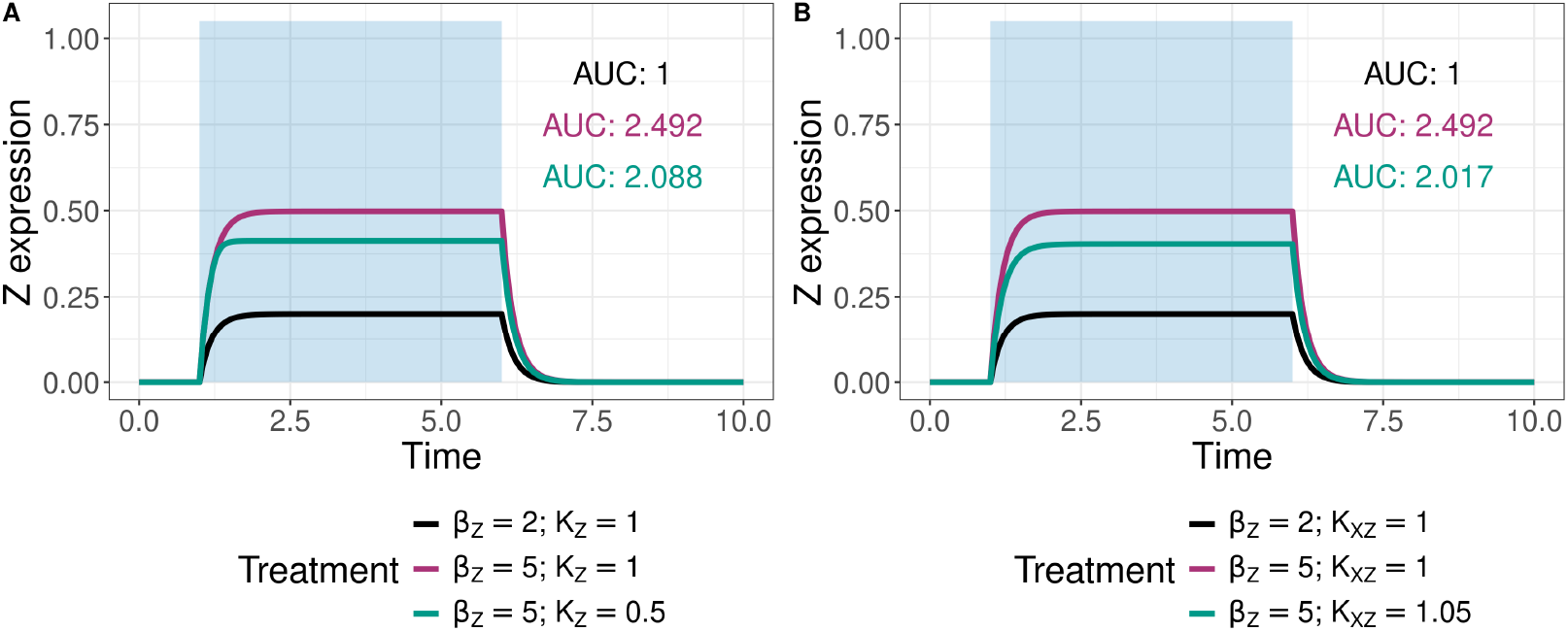
Demonstration of the interaction between *K* parameters and *β*_*Z*_ on *Z* expression. The black line shows a baseline scenario, the red line shows a mutation in *β*_*Z*_ alone creating a large effect on trait expression. In (A), the green line shows the same *β*_*Z*_ mutation with a reduced effect on *Z* expression due to an additional reduction in *K*_*Z*_. In (B), the green line shows the same scenario as in (A), but by a small increase in *K*_*XZ*_. AUC is the area under the curve (the trait value). For all curves, *α*_*Z*_ = 5. In A, *K*_*XZ*_ = 1 and in B, *K*_*Z*_ = 1.

*K*_*XZ*_ has the opposite effect to *K*_*Z*_: it controls the concentration of *X* required to produce half-maximal *Z* (Alon 2019). *K*_*XZ*_ mutations alter the amount of expression by adjusting the time for *Z* expression to reach steady-state (Rosenfeld *et al*. 2002). This can have large effects on expression by altering the time spent producing the maximum amount of *Z* (Fig. 10B). We observe the power of this adjustment by the larger fitness effects *K*_*XZ*_ mutations tend to carry (Fig. 9). Under the small effect regime, this gives K+ models an axis of molecular variation which can produce larger steps towards the optimum, especially because of their interactions with other molecular components (Fig. 3).

The *K* parameters interact with *α*_*Z*_ and *β*_*Z*_ mutations to recontextualize their fitness effects (Fig. 10). This leads to the larger fitness effects among *K*_*XZ*_ mutations we observed, however in *K*_*Z*_, such mutations are not sampled due to the small effect size treatment. Our previous work showed that *α*_*Z*_ and *β*_*Z*_ mutations in K− models could have strongly deleterious effects on fitness and that such alleles were fairly common (O’Brien *et al*. 2024). The K+ model appears to introduce complexity to this via the large beneficial fitness effects of *K*_*XZ*_ mutations relative to the other molecular components (Fig. 8, 9).

Our observation of increased variability between replicates of evolution in K+ models under low recombination lends credence to this theory (Fig. 5, 7). Given the context-dependency of the fitness effects of mutations in molecular components, selection is expected to favor the inheritance of beneficial genotypes rather than individual alleles. As a result, the molecular architecture of the trait has ramifications for the dynamics of alleles at a population scale.

### NAR-mediated linkage disequilibrium

In K+ models, we observed an excess of both positive and negative linkage disequilibrium (LD) which exceeded that of K−populations (Fig. 4). This was likely a result of the epistasis emerging from the network structure. However, on average, LD remained similar between the models, being close to 0. Given that positive epistasis should generate positive LD, this was unexpected (Otto 2009, Fig. 3).

Two mechanisms likely contribute to this pattern. First, Hill-Robertson interference can mask the relationship between epistasis and LD: when drift is strong relative to selection, beneficial alleles may be lost if they initially occur on disadvantageous backgrounds, even with just two loci. This phenomenon has been extensively documented in experimental populations of *Drosophila melanogaster*, where regions of low recombination show reduced efficacy of selection and decreased nucleotide diversity (Campos *et al*. 2014). Similarly, studies in *Saccharomyces cerevisiae* have demonstrated that beneficial mutations occurring on different genetic backgrounds compete with each other, leading to slower adaptation rates than predicted by their individual effects (Lang *et al*. 2013). These empirical observations align with our finding that K+ populations show more extreme LD patterns despite consistent positive epistasis. While Hill-Robertson interference explains the immediate effects of selection and drift on beneficial mutations, a second mechanism operating at the network level further complicates the relationship between epistasis and LD.

Second, higher-order epistatic interactions could mask the pairwise effects we measured. This is supported by experimental work in bacterial systems: measurements of higher-order epistasis in several bacterial enzymes by Buda *et al*. (2023) revealed that the majority of mutations disrupted existing epistatic networks, resulting in fluctuations between positive and negative epistasis (idiosyncratic epistasis). Similar patterns have been observed in antibiotic resistance evolution, where Weinreich *et al*. (2006) found that the fitness effects of resistance mutations were highly dependent on the order of acquisition, suggesting complex interaction networks beyond pairwise effects. However, idiosyncratic epistasis is expected to slow adaptation (Lyons *et al*. 2020), so further analysis into the strength of multilocus epistasis across molecular components would be helpful to quantify how epistasis contributes to fitness across replicates of evolution. The prevalence of such higher-order interactions in empirical systems supports the complex LD patterns we observed in the K+ model, where the more intricate network architecture provides greater potential for multilocus interactions. We also observed that while pairwise epistasis was similar between molecular components, there still was a consistent difference between the K+ and K− models, lending credence to this theory (Fig S7, Fig 3).

The observation of slightly positive fitness epistasis in the additive model 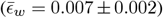 was unexpected, given the Gaussian fitness function’s negative curvature (Kingsolver *et al*. 2001) and zero trait epistasis. In theory, a Gaussian fitness function should generate negative epistasis between beneficial mutations in populations near the optimum, as the combined effect of two beneficial mutations would overshoot the optimum (Mäki-Tanila and Hill 2014). However, several factors could alter this expectation.

First, our small effect size regime (*τ* = 0.0125) meant that over-shooting the optimum with pairs of mutations was unlikely, even with positive epistasis. Second, even if overshooting occurred, the fitness cost would be minimal due to the small phenotypic effects. Third, while we measured epistasis across the entire adaptive walk, the stronger fitness gains early in adaptation (when populations were far from the optimum) might have outweighed any negative epistasis near the optimum. However, this explanation is complicated by our observation that mean fitness epistasis remained constant throughout the adaptive walk. Further work explicitly tracking the relationship between distance to the optimum and the strength of fitness epistasis would help to resolve this puzzle. While the additive model showed positive fitness epistasis, its magnitude was much smaller than in the network models, suggesting that network architecture, rather than the fitness function, was the primary driver of epistatic interactions in our simulations.

### The omnigenic model and network models of adaptation

Our attempt to reconcile quantitative genetics theory with the molecular machinery underpinning gene expression is not the first. The omnigenic model posits that large effect QTLs represent rare, highly-connected *cis*-acting elements (“core” genes) and that small effect QTLs are *trans*-acting transcription factors and other regulatory elements (“peripheral genes”) which affect core gene expression rather than the trait directly (Liu *et al*. 2019; Boyle *et al*. 2017).

In our model, QTLs affecting *K*_*XZ*_ represent core genes, whereas the other molecular components behave more similarly to peripheral genes. In the omnigenic model, core genes have large effects owing to their high network connectivity: they affect the phenotypic effects of many loci via genetic interactions (Boyle *et al*. 2017). These interactions can result in existing deleterious alleles to become neutral or beneficial. This results in a sudden increase in the frequency of those alleles and an increase in *V*_*A*_, as previously inaccessible genetic variation is released and usable for adaptation (Goodnight 1988). The network connectivity of transcription factors (or QTLs in the context of our simulations) might then hint at the ability of mutations to recontextualize alleles at other loci, or the robustness of interactions to new mutations.

We observed the consequences of the omnigenic model in our K+ populations via elevated epistasis (Fig. 3), LD (Fig. 4), and *V*_*A*_ (Fig. 2), resulting in a robustness to recombination that was not seen in the other models (Fig. 1). The interplay between the above factors creates a web of terms contributing to additive variance. Different networks and genetic architectures would be expected to produce additive variance with different combinations of epistasis, linkage, recombination, and mutational effects. Over long periods, these different combinations might produce reproductive incompatibilities via system drift.

Under system drift, neutral mutations (regarding traits under selection) drive changes in molecular architecture, which can eventually lead to reproductive incompatibilities and speciation (Schiffman and Ralph 2021). This could occur following adaptation when trait values stabilize around a phenotypic optimum and most segregating *trans-* mutations are neutral (e.g., the *K*_*Z*_ mutations in our simulations, Fig. 9). However, owing to specific changes in the contextualizing *cis-* mutations (e.g., *K*_*XZ*_ alleles), attempted crossing of *trans-* mutations between lines with different *cis-* configurations could be inviable. Prior simulation studies (for example, see Johnson and Porter 2000, 2007; Schiffman and Ralph 2021) have found system drift can arise under stabilizing selection when pleiotropy is common. Schiffman and Ralph (2021) found that genetic networks could contribute to system drift. Our results corroborate this, suggesting that network structure and linkage between molecular components might provide circumstances favoring system drift. Future work identifying these circumstances will be illuminating, as will further investigations into more complex genetic networks.

Our results highlight the fickleness of adaptation via network-mediated quantitative traits. Under larger effects, the differences between models were considerably less apparent (Fig. S2, S3). Indeed, the additive model was less affected by recombination when mutations were larger (fewer combinations were required to reach the optimum, reducing the need for recombination). Similarly, the gap between K+ and K− models also shrank with increases in effect size (Fig. S1, S2, S3). The common trend was that the fewer steps required to reach the optimum, the smaller the difference between models. With more steps, there is more opportunity for interactions to arise and accumulate, leading to the larger differences between models that we observed under the small effect regime.

Given that effect size can alter the adaptive walk, changes in the network structure are likely to produce very different evolutionary dynamics. Even relatively small changes to the NAR model, such as using a Hill function for *X* instead of a step function, would likely tune the emergence of nonlinear interactions. Previous work exploring topological features of networks on additive variance suggests that the choice of the network also plays a role in adaptation. Malcom (2011) found that network size was negatively correlated with the release of additive variance, i.e., smaller networks could release additive variance faster than larger networks. Since we consider a simple network with only (up to) four evolving components, the NAR model is firmly in the small network space where *V*_*A*_ is expected to be quickly released, which aligns with our results (Fig. 2).

In another vein, Macía *et al*. (2012) found that positive epistasis was a feature of simple networks, whilst, in complicated structures, negative epistasis was more common. In addition, robustness to mutation was common in complex networks. This suggests that recombination could be favored in more complex networks, as the breaking down of negative LD produced by negative epistasis increases additive variance in fitness (Barton 2010). Our simple NAR system harbored mostly neutral LD, but both positive and negative LD under low recombination, disagreeing with Macia et al.’s (2012) findings. In our case, the genetic architecture, particularly the emergence of Hill-Robertson effects, might have played a more important role than network complexity.

Our results indicate that both the genomic distribution of allelic variation and the underlying networks describing the shape and size of that variation are key for determining the role of the genotype-phenotype map in shaping quantitative traits. Future work exploring how different network structures might mediate *V*_*A*_ would be helpful in ascertaining the general importance of network-derived epistasis to *V*_*A*_ and the rate at which contextualizing mutations can spread through populations. It will also reveal how additive variance is distributed within quantitative traits, how network structures contribute to the standing variation required for adaptation, and how recombination contributes to additive variance under different genetic architectures.

### Limitations and Assumptions

Our study provides valuable insights into how genetic network structures influence polygenic adaptation, but we must acknowledge certain limitations and assumptions inherent in our models. One key assumption is our use of a step function to model *X* expression in the NAR network (Eq. 2). This step function represents an instantaneous activation of *X* in response to an environmental cue, simplifying the dynamic nature of gene expression. In biological systems, gene activation often follows a more gradual response, which can be modeled using a sigmoid curve to reflect cooperative binding and threshold effects (Alon 2019; Hill 1910). The abrupt activation in our model may influence the timing and levels of *Z* expression, potentially affecting the strength and nature of the genetic interactions observed. Future studies could explore alternative activation functions for *X* to assess the robustness of our findings to changes in gene activation dynamics.

We also chose to sample mutational effects from a normal distribution. This is a common assumption in studies of quantitative traits (e.g., Lande (1976)). However, in natural populations, the distribution of mutational effects on quantitative traits can be leptokurtic or long-tailed, meaning that large effects are sampled more frequently than under a normal distribution (Mackay and Lyman 1998; Schraiber and Landis 2015). While our chosen *τ* values enabled us to capture more frequent large effects, they may not precisely align with empirical distributions observed in specific organisms or traits. Incorporating empirical data on mutation effect sizes and recombination rates would enhance the biological realism of our models and the applicability of our findings to natural populations.

Additionally, certain parameters in our ordinary differential equations (ODEs) were fixed to balance biological realism with computational tractability. For instance, the Hill coefficient *h* was set to 8 to produce a steep activation curve, mirroring strong cooperative binding in some gene regulatory systems. However, variations in *h* could alter the sensitivity of the system to changes in molecular components, potentially affecting the emergence of epistasis and the distribution of additive variance. Similarly, the time boundaries for *X* activation (*t*_start_ = 1, *t*_stop_ = 6) were chosen arbitrarily to represent a specific environmental exposure period. Altering these time frames could influence the total *Z* expression and its response to selection. While our parameter choices capture essential features of the NAR network, exploring a wider range of parameter values would enhance our understanding of how these assumptions impact adaptive dynamics. In a similar vein, we chose our recombination rate treatments *a priori*; however, there is no guarantee that all of these rates would be able to arise or persist in natural populations, especially on a genome-wide scale.

Our analysis focused on estimating pairwise epistasis between segregating alleles to understand how genetic interactions contribute to adaptation. While pairwise epistasis provides valuable insights into the nature and direction of genetic interactions, it does not capture the full complexity of multilocus interactions that can occur in biological systems (Weinreich *et al*. 2013). Higher-order epistasis, involving interactions among three or more loci, can significantly influence evolutionary trajectories by affecting the shape of the fitness landscape and the accessibility of adaptive paths (Weinreich *et al*. 2013; Sailer and Harms 2017).

Multilocus interactions are likely in complex genetic networks like the NAR motif, given the interconnectedness of molecular components and the potential for emergent properties arising from network dynamics. While measuring higher-order epistasis poses computational challenges, insights into three-way or four-way interactions might be sufficient for capturing much of the variance in fitness effects (Weinreich *et al*. 2018). Future work could employ methods such as Fourier-Walsh transformations (used in Weinreich *et al*. (2018) for example) to interrogate the importance of higher-order interactions arising from genetic networks on the adaptive landscape and the maintenance of genetic variation in natural populations.

## Conclusions

Our findings suggest that the underlying mechanisms of traits can direct the adaptive outcomes of populations. The robustness of the K+ model to low recombination rates indicates that organisms with complex gene regulatory networks may better adapt to environmental changes, even under limited recombination. This is especially relevant for species with small effective population sizes, or populations in regions with suppressed recombination due to chromosomal inversions or structural genomic features. Understanding how network complexity influences genetic variation and epistasis is crucial for predicting evolutionary trajectories.

Rapid adaptation is vital for survival in species facing climate change, habitat fragmentation, or novel pathogens. Our study highlights the importance of molecular architecture in assessing adaptive potential, as genetic network structures can greatly influence additive genetic variance and selection efficiency. Future work exploring how these mutations influence system drift in simple and complex networks could clarify the connection between adaptation, speciation, and the distribution of additive and nonadditive variation in quantitative traits.

## Author contributions

DO conceived the original idea. NO, JE, BH, and DO jointly developed the model and designed simulation experiments. NO implemented and ran the simulations and analysis. BH and JE provided mathematical guidance. NO wrote the paper with assistance from all authors.

## Data availability

The data generated in this study is available at: https://doi.org/10.48610/1a19d80. Scripts to run the analysis are available at: https://github.com/nobrien97/PolygenicNAR2024 and SLiM scripts to run the simulations. SLiM modifications are available at https://github.com/nobrien97/SLiM/releases/tag/PolygenicNAR2024: note that this modification requires the Ascent ODE solution library installed https://github.com/AnyarInc/Ascent/releases/tag/v0.7.0.

## Acknowledgments

We acknowledge the Turrbal and Yuggera people, traditional owners of the land on which this work was undertaken, and pay their respects to Elders past, present, and emerging. We would like to acknowledge members of the Ortiz-Barrientos lab for their feedback on this research. This research was carried out using the Gadi HPC system maintained by the National Computational Infrastructure (NCI), which is supported by the Australian Government.

## Supplementary material

### S1 Appendix

We chose the *τ* levels based on a basic estimate of the expected number of beneficial mutations to take a population to the optimum. Mutational effects are sampled from a normal distribution, *a* ~ 𝒩 (0, *τ*), where *a* is the mutational effect on the trait (additive) or the molecular component (network). Assuming that adaptation is driven entirely by beneficial mutations (i.e., all mutations drive the population closer to the optimum), the distribution of adaptive mutations is given by a folded normal. The folded normal has expected value

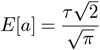

where *E*[*a*] is the expected effect size for *a*. Rearranging, we can find the *τ* required to produce a given *E*[*a*]. Further we can find the number of additive mutations required to reach a given *E*[*a*] by introducing another term, *n*:

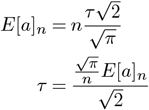

The total phenotypic change required to reach the optimum is one, which we can substitute into *E*[*a*]_*n*_. Then, we can estimate the *τ* required to reach the optimum in *n* beneficial mutations. Using this method, we calculated *τ* = 1.25 corresponds to 1 mutation required to reach the optimum, *τ* = 0.125 for ten mutations, and *τ* = 0.0125 for 100 mutations required. While this is a crude approach (which ignores the epistatic effects expected in the network models and the likelihood of compensatory mutations), it gives a rough quantification of “large”, “intermediate”, and “small” effects in our study.

**Supplementary Figure 1.**
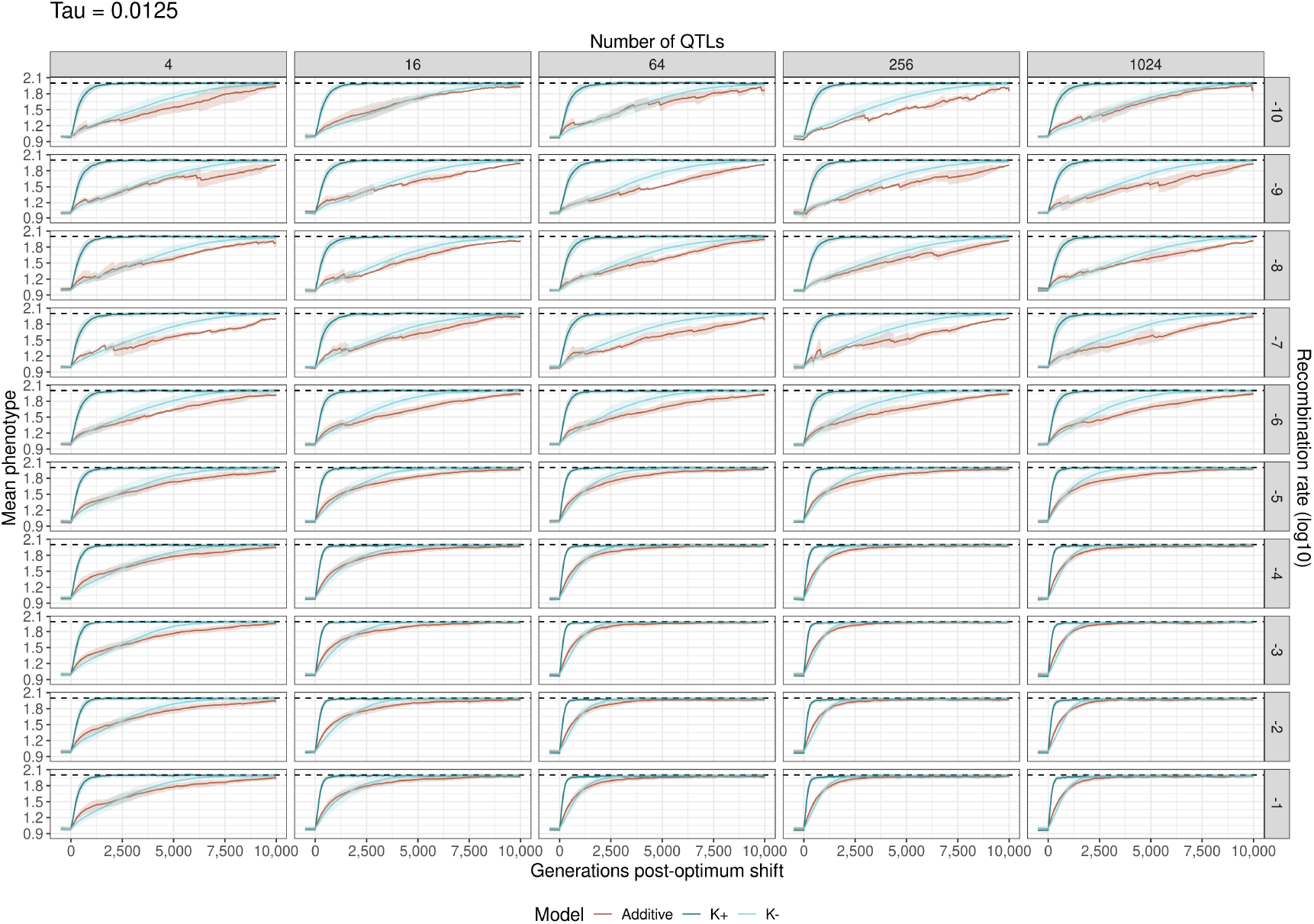
Adaptation to a shifting optimum in all tested number of loci/recombination rate treatments and under small mutational effects (*τ* = 0.0125). Each line is the mean of population among populations that reached the optimum within 10,000 generations. There were up to 48 replicates per group. Ribbons show 95% confidence intervals. The dashed line shows the phenotypic optimum at *O* = 2.

**Supplementary Figure 2.**
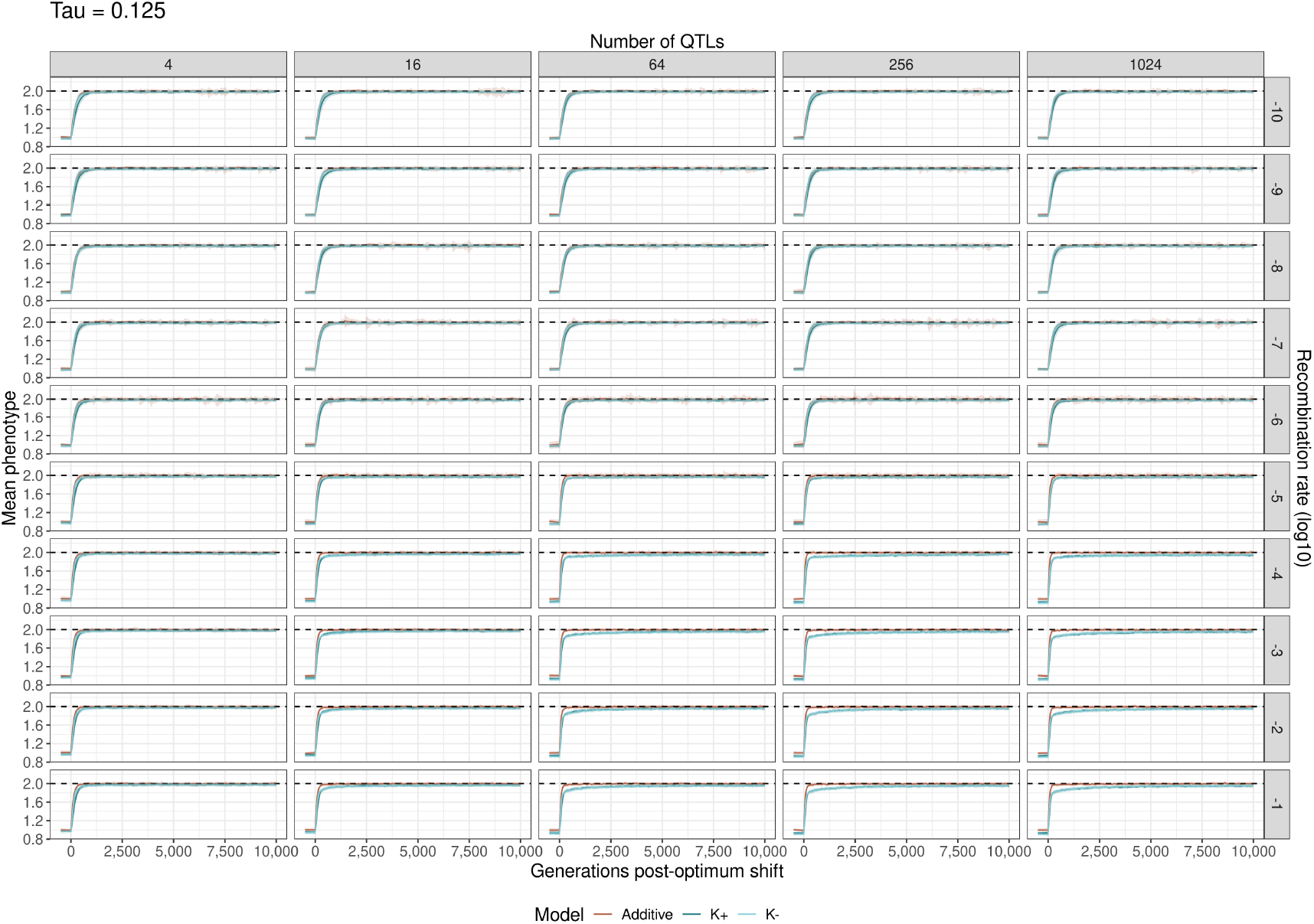
Adaptation to a shifting optimum in all tested number of loci/recombination rate treatments and under intermediate mutational effects (*τ* = 0.125). Each line is the mean of population among populations that reached the optimum within 10,000 generations. There were up to 48 replicates per group. Ribbons show 95% confidence intervals. The dashed line shows the phenotypic optimum at *O* = 2.

**Supplementary Figure 3.**
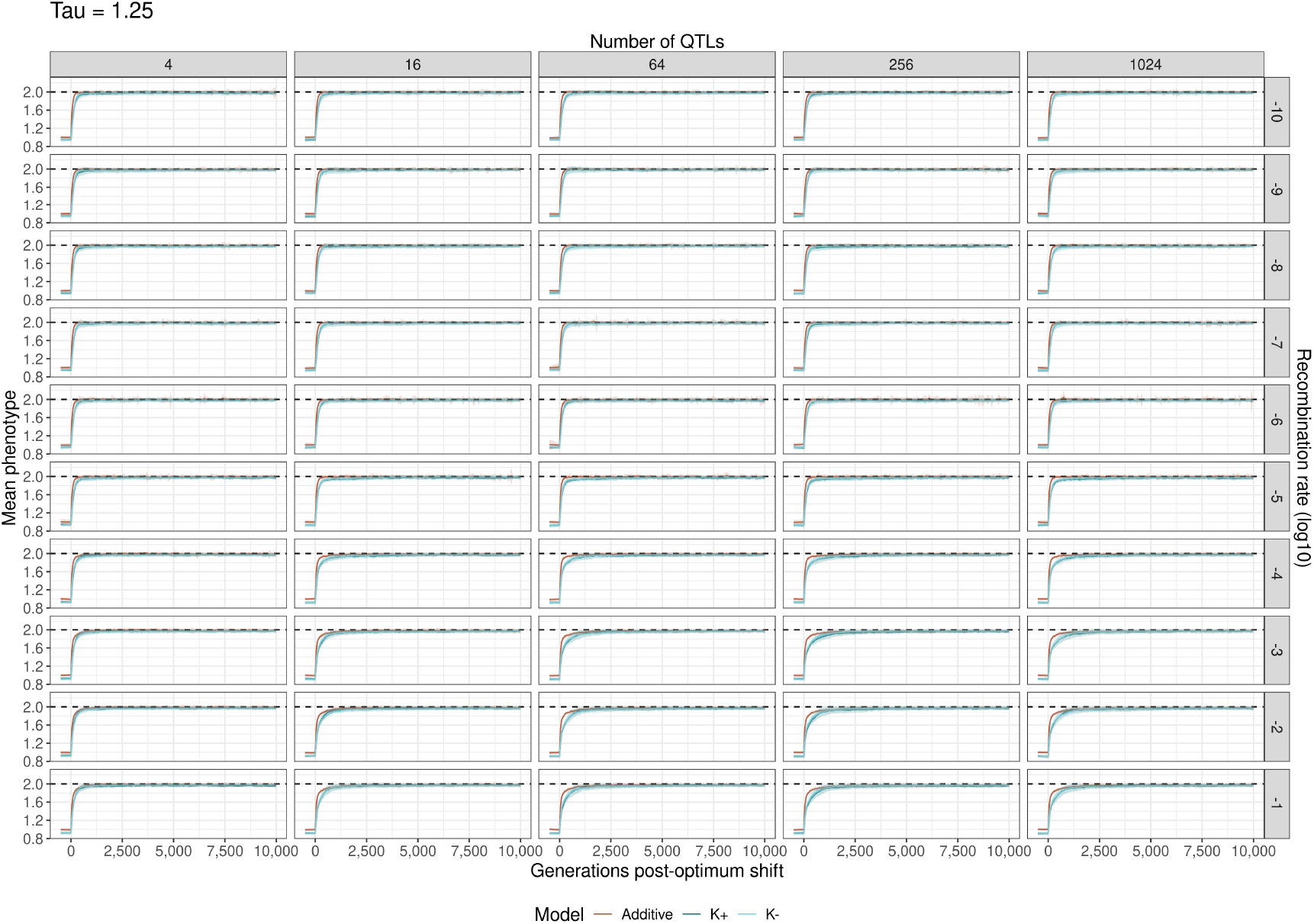
Adaptation to a shifting optimum in all tested number of loci/recombination rate treatments and under large mutational effects (*τ* = 1.25). Each line is the mean of population among populations that reached the optimum within 10,000 generations. There were up to 48 replicates per group. Ribbons show 95% confidence intervals. The dashed line shows the phenotypic optimum at *O* = 2.

**Supplementary Figure 4.**
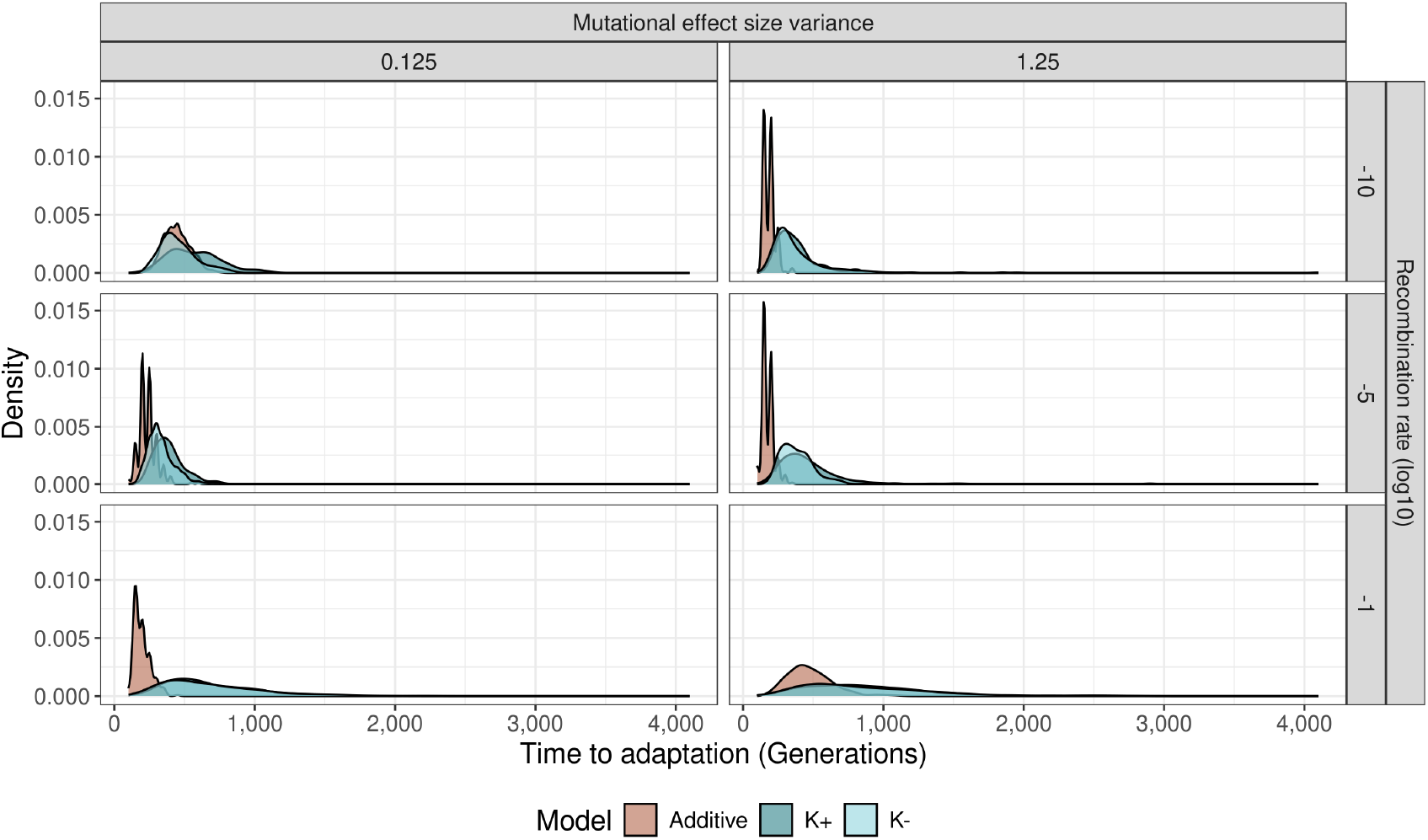
Number of generations to reach the new phenotypic optimum among all models and replicates with intermediate and large mutational effect size variance treatments. Densities are measured across the five number of loci treatments.

**Supplementary Figure 5.**
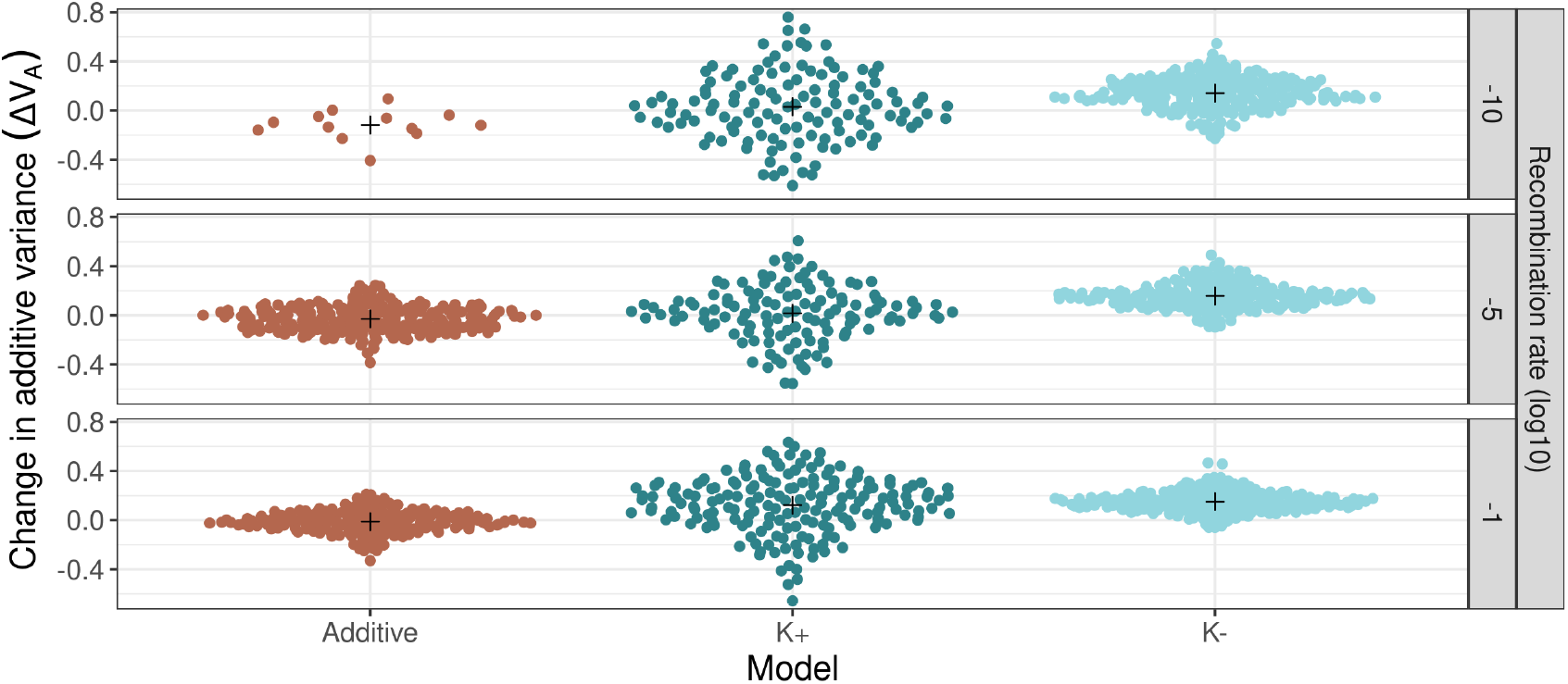
Change in additive variance (Δ*V*_*A*_) during adaptive walks for each model and recombination rate. Points represent populations, and crosses represent group means. Δ*V*_*A*_ is measured across the entire adaptive walk.

**Supplementary Figure 6.**
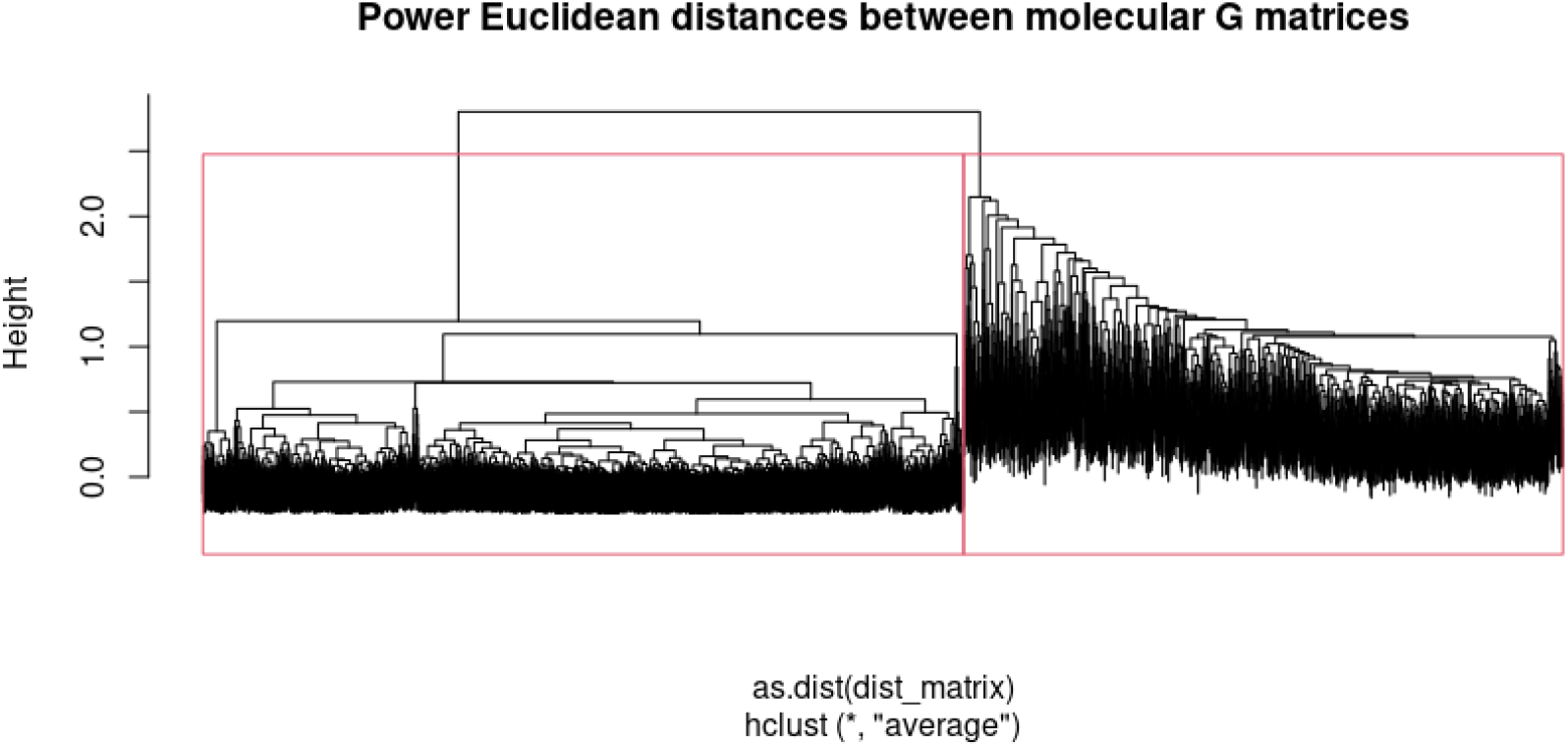
Dendrogram of hierarchical clustering analysis of Power Euclidean distances between **G**_*C*_ matrices. Red boxes show chosen clusters.

**Supplementary Figure 7.**
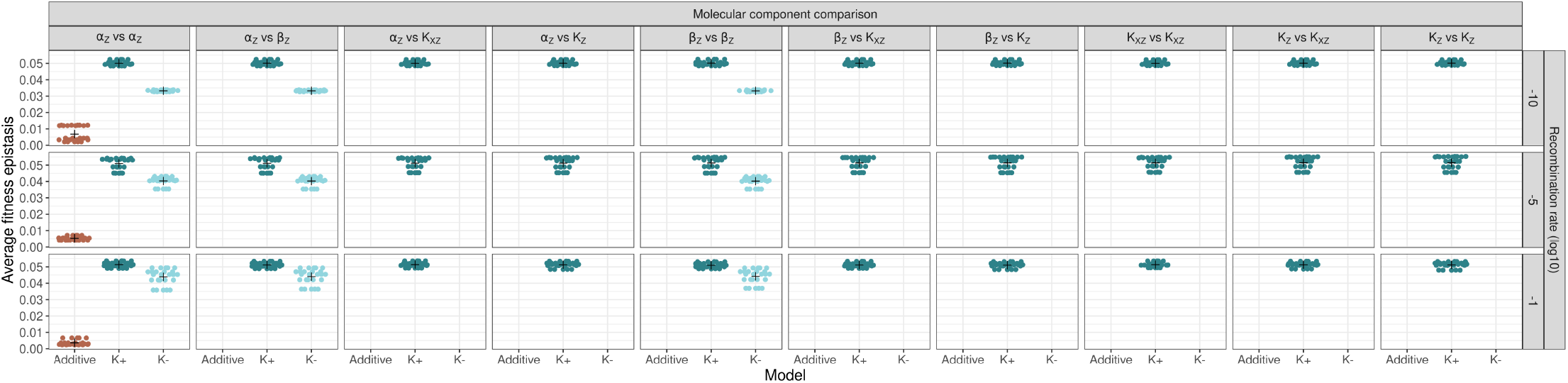
Mean pairwise fitness epistasis 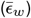 in adapted populations with small mutational effect size variance (*τ* = 0.0125) across each pairwise comparison of molecular components. Reciprocal comparisons (e.g., *α*_*Z*_ vs *β*_*Z*_ and *β*_*Z*_ vs *α*_*Z*_) are averaged. Each point represents 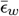 in a single population. The cross represents the mean 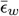 across different numbers of loci treatments and time points.

